# The Caudate Nucleus Exhibits Distinct Pathology and Cell Type-Specific Responses Across Alzheimer’s Disease

**DOI:** 10.64898/2026.01.10.694705

**Authors:** Omar Z. Kana, Nadia Postupna, Anamika Agrawal, Brian Long, Kyle J. Travaglini, Mariano I. Gabitto, Hsin-Yu Lai, Joseph T. Mahoney, Ming Xiao, Tejas Bajwa, Heino Hulsey-Vincent, Sora L Oyaizu, Gonzalo E. Mena, Janna Hong, Emily C. Gelfand, Eitan S. Kaplan, Jeanelle Ariza, Kimberly A. Smith, Delissa A. McMillen, Samantha D. Hasting, Anish Bhaswanth Chakka, Jeff Goldy, Erica J. Melief, Zoe Juneau, Stuard Barta, Alana Oyama, Augustin Ruiz, Christina Alice Pom, Angela Ayala, Madeline Bixby, Alvin Huang, Noami Martin, Nasmil Valera Cuevas, Paul Olsen, Josh Nagra, Rushil Chakrabarty, Michael Tieu, Trangthanh Cardenas, Amy Torkelson, Darren Bertagnolli, Junitta Guzman, Rebecca Ferrer, Jack Waters, Thomas J. Grabowski, Paul K. Crane, Nicole M. Gatto, Jeremy A. Miller, Rebecca D. Hodge, Michael Hawrylycz, C. Dirk Keene, Ed S. Lein, Jennie L. Close

## Abstract

Aβ presence in the caudate nucleus (Ca) partially defines Thal stage III in Alzheimer’s disease (AD), but little is known about AD’s cellular impact on the region. Leveraging a public basal ganglia taxonomy of cellular populations, we generated a cellular resolution atlas of AD-associated pathological changes in Ca. Unlike cortex, we found that Ca AD pathology is dominated by two key features: phosphorylated tau (pTau)-containing neuropil threads enriched near oligodendrocytes in white matter tracts and amyloid-β diffuse plaques enriched in gray matter. Although AD pathology in affected cortical regions results in neuronal loss, we find no AD-driven reductions in neuron proportions in Ca. However, there were observable changes in multiple cellular populations. Protoplasmic astrocytes and FLT1+/IL1B+ microglia increased in abundance with global pTau levels. We also observe gene expression changes in fast-spiking PTHLH-PVALB interneurons indicative of disrupted signaling pathways and altered intrinsic physiological properties. This work provides a cellular-resolution framework for understanding AD pathology in Ca.

## INTRODUCTION

The Brain Initiative Cell Atlas Network (BICAN) has recently employed single cell transcriptomics and spatial methods to characterize cellular gene expression and spatial relationships in the brains of multiple species at high resolution^1–4^. In parallel, the Seattle Alzheimer’s Disease Brain Cell Atlas consortium (SEA-AD), has adopted this paradigm to study the cellular and molecular changes that occur during Alzheimer’s disease(AD)^5,6^. BICAN’s most recent effort culminated in a standardized and reproducible community taxonomy that integrates existing atlases to describe the human basal ganglia (BG) and its unique cell types^2,7–9^. The BG are made up of several regions including, but not limited to, the globus pallidus, the substantia nigra, the putamen, and the caudate nucleus (Ca). The Ca, as part of the cortico-basal ganglia-thalamic loop, receives excitatory input from the cortex and thalamus, as well as dopaminergic inputs from the substantia nigra pars compacta, and exerts inhibitory output on the substantia nigra pars reticula and the globus pallidus (Figure 1A)^10^. This implicates the Ca in many brain functions including learning new motor skills and selecting contextually appropriate actions^11^. Dysfunction of the Ca is at the nexus of neurodegenerative diseases such as Huntington’s disease and Parkinson’s disease and contributes to several symptoms such as cognitive impairment, depression, disordered sleep and impaired movement (Figure 1B) ^12–23^. In Huntington’s, this may be connected to a loss of Ca medium spiny neurons (MSNs)^24^. In Parkinson’s, MSNs exhibit more physiological changes, such as a reduced spininess characteristic^25,26^. MSNs affected by Parkinson’s also display reduced dopaminergic connections and dopaminergic response^27–29^. AD can be comorbid with both these diseases, making it difficult to disentangle the effects of one disease from another^30–33^.

**Figure 1.**
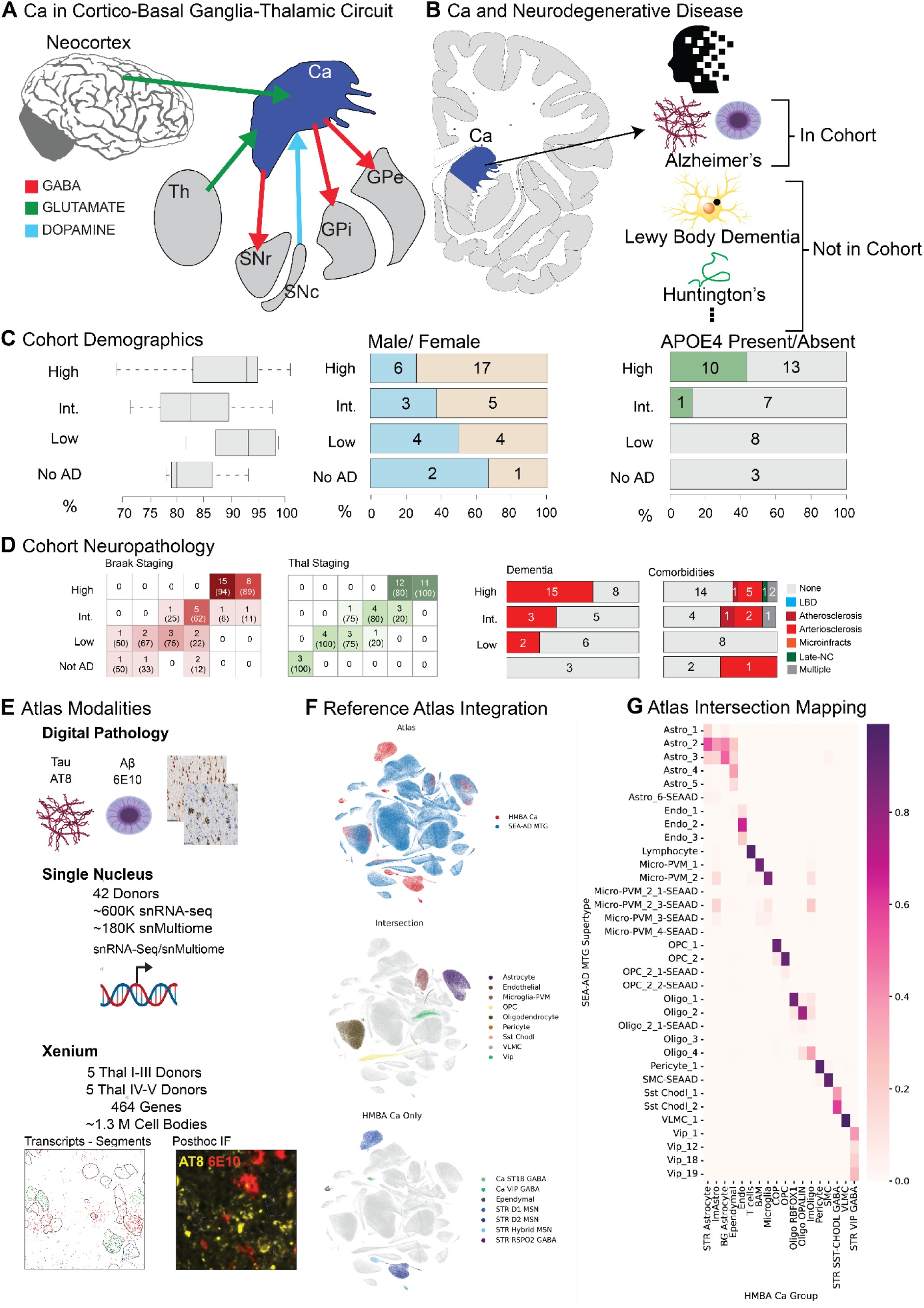
A molecular atlas of the Ca in AD. (A) Schematic of inputs and outputs to Caudate nucleus (Ca), which receives glutamatergic (green) input from the cortex and thalamus (Th), dopaminergic input (blue) from substantia nigra pars compacta (SNc), and sends GABAergic output (red) to substantia nigra pars reticulata (SNr), and Globus pallidus externus (GPe) and internus (GPi) (B) Sampling of the Ca and its associated neurodegenerative pathologies. Alzheimer’s pathologies are enriched in the cohort; comorbid diseases are minimized. (C) SEA-AD cohort demographics, depicting age at death (left, box represents (IQR), whiskers represent 1.5 times the IQR, median indicated by solid line), biological sex (middle) and APOE4 allele status (right), stratified according to ADNC score. (D) SEA-AD cohort composition stratified according to ADNC versus Braak stage and Thal phase (left), with dementia or comorbidities as bar plots (right). Donor number is indicated in each box, fraction shown in parentheses. (E) Illustration of molecular assays (snRNA-seq/snMultiome, and Xenium spatial transcriptomics) and neuropathology applied to AD Spectrum of SEA-AD cohort. (F) UMAP coordinates of cellular latent space of SEA-AD middle temporal gyrus (MTG) atlas and HMBA basal ganglia (BG) atlas subsetted to caudate nucleus (Ca) produced by scANVI. Top: colored by atlas, Middle: colored by HMBA BG atlas exclusive cell types, Bottom: colored by cell types in both atlases. (G) Mapping between cell type labels in HMBA BG atlas to SEA-AD MTG atlas. All columns are normalized by total number of cells in each HMBA BG atlas label.

AD, as defined by accumulation of amyloid beta plaques (A*β*) in the cortex and phosphorylated tau (pTau) tangles, begins in the entorhinal cortex and involves the hippocampus early in disease progression^34,35^. Previous efforts have shown loss of inhibitory interneurons, excitatory intratelencephalic neurons, and oligodendrocyte populations, along with increases in microglia and astrocyte populations with Alzheimer’s in cortical areas ^5,36,37^. However, AD pathology eventually spreads to every cortical region as well as many subcortical regions in surviving affected individuals. The Ca is affected by AD neuropathology, albeit usually in individuals with already high levels of cortical pathological burden^38,39^. Additionally, the Ca is known to atrophy in direct proportion with AD pathology burden as the disease progresses ^34,40–44^. Ca dysfunction may also contribute to the cognitive and behavioral changes observed in AD^45,46^. In a study of the ROS/MAP cohort, bulk expression of VEGF signaling in the Ca was linked to neuroprotective effects in AD pathological progression^47^. In addition, a recent large multiregional atlas study linked increased neuritic plaque burden in the prefrontal cortex to increases in SERPINH1+ caudate astrocytes while mild cognitive impairment was linked to increases in WIF1+/COX8A+ astrocytes in Ca. In that same study, expression of SLC5A3 in oligodendrocytes was correlated with several pathological features of AD^22^. However, the authors note that this study did not use quantitative neuropathological information specific to the Ca region beyond presence and absence of A<u>β</u> plaques for Thal staging^48^. In another study, microglial responses to pathology in caudate appear to differ from those of their cortical counterparts in that activation of these cells does not include known neuroinflammatory responses in this context^49^. In addition, there may be differences in the nature and distribution of pathology in caudate given the distinct cellular composition and processing demands of the basal ganglia.

Despite these initial studies and the central role of the basal ganglia in decision making, motor control and reward processing, the cellular and molecular significance and impact of AD pathology in the caudate nucleus remains unclear. We took advantage of the novel information regarding caudate cell types to generate a multi-modal atlas of the caudate nucleus of 42 individuals spanning Braak and Thal staging, and determine the impact of AD progression on the gene expression programs of caudate cellular populations ^2,7,8^. Quantitative neuropathology was performed on the same donors to provide context for our cellular observations. We also selected a subset of 10 donors for spatial transcriptomics profiling with post-hoc staining for Aβ and pTau to determine local effects of pathology on gene expression and cellular response. Through quantitative neuropathology we find that diffuse Aβ plaques and tau neuropil threads constitute most of the AD pathology in Ca in these donors. We observe that the neuropil threads tend to co-localize with oligodendrocyte-rich regions or white matter and that the diffuse plaques are most often localized to grey matter. Unlike AD-affected cortical regions or basal ganglia impacted by Huntington’s disease, there are no significant changes in the abundance of neuronal types with AD progression in Ca. However, we do observe an increase in the abundance of protoplasmic astrocytes and FLT1+/IL1B+ microglia in this region. Though there were no neuronal abundance changes, fast-spiking PTHLH-PVALB interneurons exhibit gene expression changes indicative of signaling disruption and altered intrinsic physiological properties. Taken together, our findings show that the response to AD pathology in Ca differs from that of affected cortical regions in these donors. This could be a result of a differing cellular milieu containing predominately GABAergic neurons, muted response to early AD-like pathology in this late-affected region, a differential make up of AD pathology compared to cortex, and/or differential compartmentalization of Aβ and pTau.

## RESULTS

### A molecular atlas of the Ca in AD

We utilized a 42-donor subset of the original 84 donor SEA-AD cohort (Figure 1C, Supplemental Table 8)^5^, minimizing comorbidities to better profile the Ca in typical AD (see STAR Methods). There were no cases of Lewy Body Dementia (LBD) and only 3 donors have limbic-predominant age-related TDP-43 encephalopathy (LATE) stage 3 and 13/42 donors have vascular comorbidities, which is common in donors of this age. High Alzheimer’s disease neuropathologic change (ADNC) donors represented 15/23 of demented individuals in the cohort and 10/23 high ADNC donors had at least one APOE4 allele. More than half of all donors were high ADNC and were Braak V or VI and Thal IV-V (Figure 1D). Females made up 5/8 of intermediate and 17/23 high ADNC donors, as expected given the higher prevalence of AD in women^34^. Donor age at death ranged from 68 years to 100 years, which is a typical range for AD related deaths.

Our atlas included three data modalities: 1) digital pathology 2) single nucleus RNA sequencing (snRNA-seq), and 3) spatial transcriptomics (Figure 1E). Digital pathology was performed on 5 µm FFPE sections from one hemisphere of each donor and included immunohistochemistry using the AT8 and 6E10 antibodies against pTau and amyloid beta A*β*, respectively. Machine learning classifiers and built-in algorithms provided by HALO software were used to quantify AT8 and 6E10 signal in each donor. Single nucleus sequencing and spatial transcriptomics were performed using adjacent fresh-frozen tissue blocks from the same hemisphere selected to include caudate head or anterior caudate body. sn RNA-seq data was generated using 10x snRNA-seq and snMultiome protocols across all 42 donors. A subset of 10 donors with few to no comorbidities was used for spatial transcriptomics. Five donors were selected for spatial data collection from the high pathology group (Thal IV-V) and five from the low pathology group (Thal I-III). Spatial transcriptomics was performed using the 10x Genomics Xenium platform with post-hoc immunofluorescence using AT8 and 6E10 antibodies.

To allow for direct comparison with previous AD studies across forebrain regions and to annotate unique cell types in the Ca, we integrated the SEA-AD middle temporal gyrus (MTG) taxonomy with the BICAN’s Human and Mammalian Brain Cell Atlas’s (HMBA) cross-species consensus cell type atlas of the basal ganglia(Figure 1F) ^2,7,8^. This allowed us to annotate known vulnerable populations in the caudate such as medium spiny neurons in Huntington’s, and specific AD associated glial states, such as microglia connected to AD progression^24,35–37^. Integration was done utilizing scANVI applied hierarchically across all levels of the SEA-AD taxonomy^38^. Except for ependymal cells, all non-neuronal populations were well integrated and thus were labeled using the existing SEA-AD taxonomy (Figure 1F, G). We observed that SST+ CHODL+ and a subset of VIP+ inhibitory interneuron populations were shared across taxonomies. The remaining neuronal populations in Ca consist primarily of lateral ganglionic eminence (LGE)-derived medium spiny neurons labeled by the Consensus BG taxonomy. This integration suggests high similarity between most caudate non-neuronal cell types with that of other cortical regions, but most of the medial ganglionic eminence (MGE) and caudal ganglionic eminence (CGE)-derived caudate neuronal populations are transcriptomically distinct from those characterized in cortical regions. The harmonized reference was used for annotation and integration with the snRNA-seq data and the total combined snRNA-seq transcriptomic reference was used for annotation of the spatial transcriptomics dataset (see STAR Methods).

### AD pathology is less pronounced in Ca compared to typical AD-affected cortical regions

We utilized conventional stains for AD staging, 6E10 and AT8, to assess Aβ plaque and pTau burden respectively (Figure 2, Supplemental Table 2). In terms of A*β*, Diffuse, compact and dense core plaques species were observed in MTG (Figure 2A-D). 6E10 signal in MTG was correlated to 6E10 signal in Ca (Figure 2I). However, in Ca we observed much lower density of dense core plaques compared to MTG, consistent with previous neuropathological studies (Figure 2E-H,J)^39–41^. 6E10 positive staining in Ca was mostly limited to diffuse plaques. Diffuse plaques have been associated with lower pathogenicity and toxicity compared to compact or dense core, suggesting that cells in the Ca could be less affected by A*β* pathology than in MTG^42,43^.

**Figure 2.**
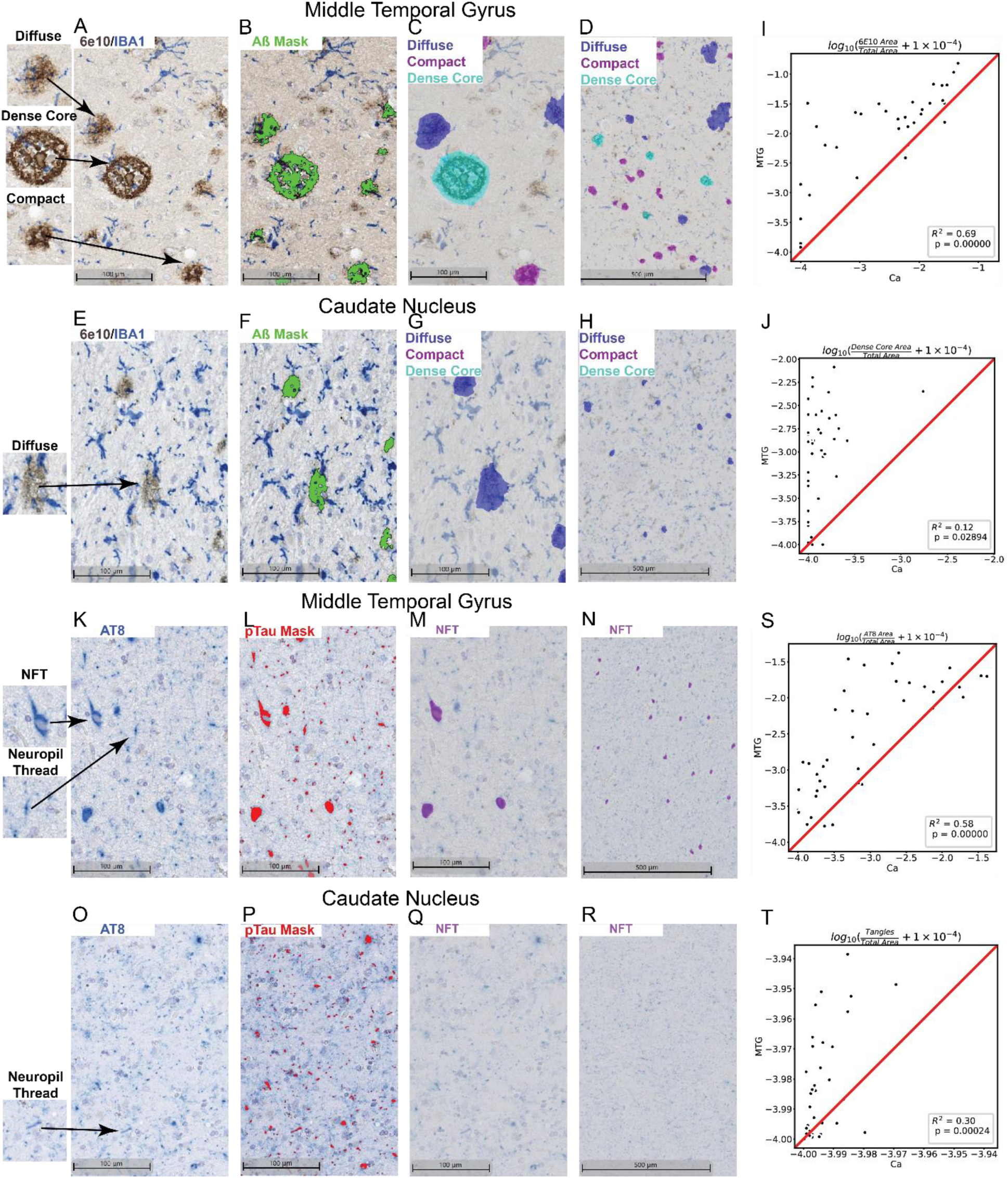
AD pathology is less pronounced in Ca compared to typical AD-affected cortical regions. (A-D) Amyloid pathology visualized with Aβ antibody (clone 6E10) in MTG and (E-H) Caudate Nucleus in donor H21.33.002. (A, E) Unmasked stain of 6E10 (brown), and IBA1 (blue). (B, F) Masked stain of 6E10 positive regions (green). (C, G) Masked regions colored by classified pathology: dense core plaques (light blue), compact plaques (magenta), and diffuse plaques (dark blue). (D, H) Zoomed out versions of (C, G). (I, J) Regression plots of pathology in MTG vs. Ca. Red line represents *y* = *x*. Comparing overall 6E10 signal in (I) and dense core plaque signal in (J). (K-N) AT8 pathological stain of pTau in MTG and (O-R) Caudate Nucleus in donor H21.33.002. (I, M) Unmasked stain of AT8 (blue). (J, N) Masked stain of AT8 positive regions (red). (K, O) Masked regions classified as neurofibrillary tangles (NFT,purple). (L, P) Zoomed out versions of (K, O). (S, T) Regression plots of pathology in MTG vs. Ca. Comparing overall AT8 signal in (S) and NFT signal in (T).

Likewise, we looked at AT8 staining in MTG and Ca to compare the appearance and extent of pTau load in these two regions (Figure 2K-T). In MTG, two patterns of pTau deposition can be observed – neurofibrillary tangles and neuropil threads (Figure 2K-N). The two pTau patterns differ in that neurofibrillary tangles are typically intracellular, forming perinuclearly in neuronal cell bodies, and neuropil threads appear in processes outside of neuronal cell bodies (Figure 2K). We note that tangles but not neuropil threads are used in diagnostic criteria of AD (as the basis for Braak staging system)^44^. Both patterns of tau appear in MTG. However, while pTau is present in Ca, qualitatively it appears predominately as neuropil threads (Figure 2O). Very few tangles were present in this region (Figure 2Q, R). Accordingly, while there is a relatively high correlation between MTG and Ca in terms of total AT8 area overall (Figure 2), neurofibrillary tangle area is much lower in Ca (Figure 2S,T). Taken together, our assessment of Ca in donors affected by AD show that this region lags behind affected cortical tissue in terms of pathological burden.

To assess the potential impact of LATE pathology, which is comorbid with AD, we stained for TDP-43 in the region. Despite LATE stage 3 donors in the study, we do not find donors with large amounts (>0.001% area) of TDP-43 in Ca (Supplemental Table 2). This suggests that LATE pathology likely has little impact on the region if any.

### Astrocyte and microglia populations increase with respect to pTau burden and Ca neurons do not change with AD progression

In MTG, increases in AD pathological burden correspond to increases in protoplasmic astrocytes and disease associated microglia, and decreases in oligodendrocytes, upper layer excitatory neurons, and SST+, PVALB+ and VIP+ inhibitory interneurons^5^. In Ca, AD pathological burden is associated with a smaller Ca volume and Huntington’s disease progression is associated with a loss of striatal MSNs^12,13,45–48^. To assess whether MTG-like AD associated cellular dynamics are present in the Ca, whether the loss of volume could be attributed to a loss in neuronal populations, or if Huntington’s neurodegeneration could be attributed, in part, to AD pathology, we model AD disease severity with an enhancement of the B-BIND algorithm^49^. Along with modeling a global donor continuous Pseudo-progression Score (CPS) based on combined AT8 and 6E10 features, to assess the specific contributions of each AD pathology we also derive feature specific (AT8 and 6E10) CPS’s (Figure 3A). We show CPS is correlated with different AD staging criteria such as ADNC (*R*^2^ = 0.85), Braak (*R*^2^= 0.55), and Thal (pearson *R*^2^ = 0.86) (Figure 3B). Global CPS is more highly correlated with 6E10 (*R*^2^= 0.58) signal than AT8 (*R*^2^= 0.37). We also observe that 6E10 pathology appears before AT8 pathology along the CPS axis (Figure 3B). Derived CPS_AT8_ for AT8 (Figure 3C) and CPS_6E10_ for 6E10 (Figure 3D) are more correlated with the progression of their respective pathologies.

**Figure 3.**
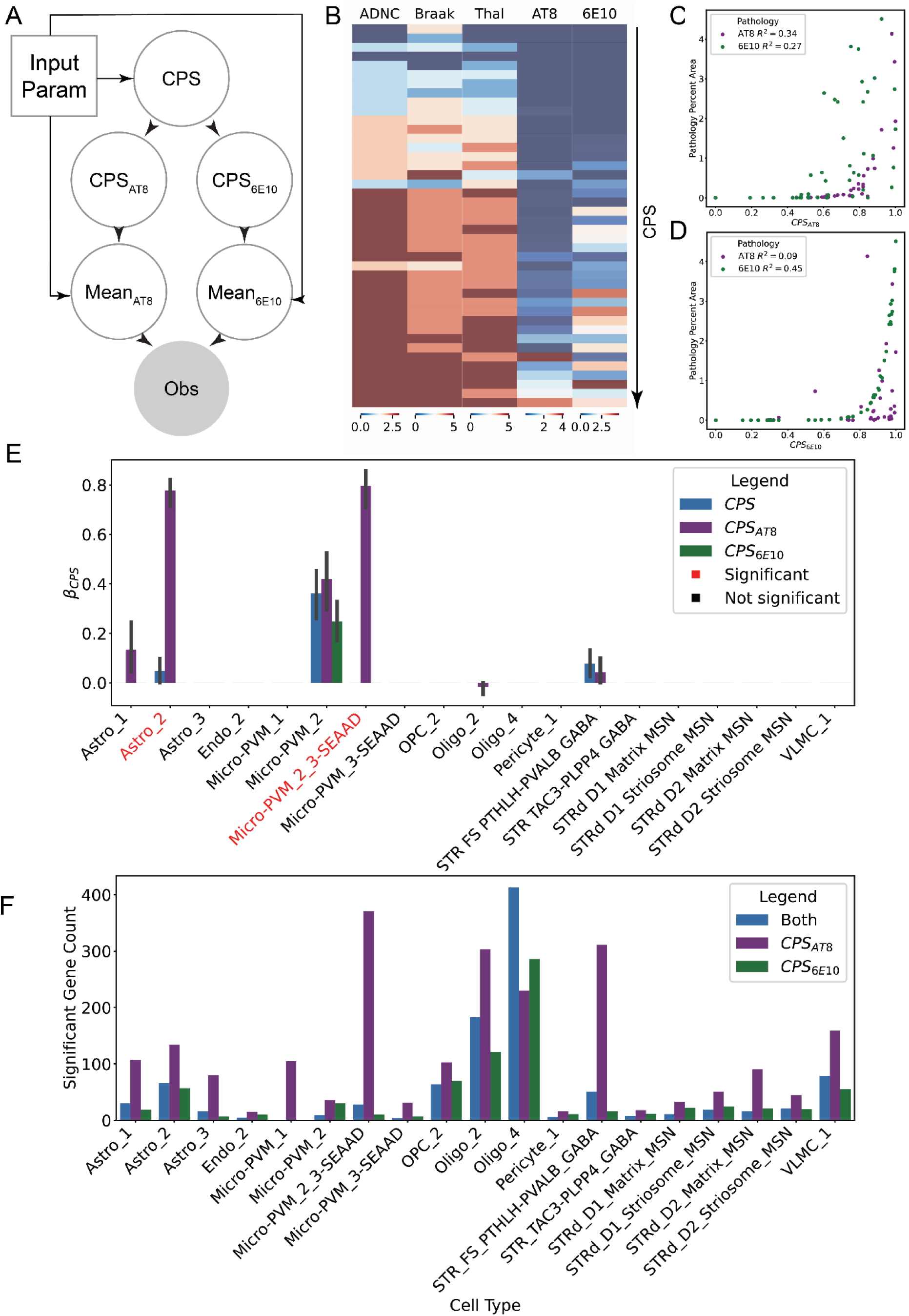
Protoplasmic astrocytes and *IL1B*+/*FLT1*+ microglia are enriched in donors with high pTau load in Ca. (A) Schematic of pathological modeling for continuous pseudo-progression scores (CPS). (B) Heatmap ordered by CPS illustrating different neuropathological scales and digital pathology in Ca. (C, D) Scatterplots visualizing the relationship between CPS_AT8_ in (C) and CPS_6E10_ in (D) versus digital pathology measurements. (E) Bar plot of *β*-coefficients from scCODA differential abundance model for global, 6E10, and AT8 CPS. Model equation: *y* ∼ *Library Prep* + *Sex* + *Age* + *APOE*4 *Status* + *CPS*_*Pat*ℎ_. Error bars represent 95% confidence interval of *β*-coefficient values across reference cell types. Significance, represented by red ticks, was defined at *p*(*inclusion*) > 0.85. (F) Bar plot of number of significantly differentially expressed genes (*p*_*adj*_ < 0.1; *β*_*CPSx*_ > 0.5) per cell type as determined by NEBULA. Model equation: *y* ∼ *Donor* + *Age* + *Sex* + *APOE*4 *Status* + 10*X Chemistry* + *CPS*_*x*_.

We mapped our Ca snRNA-seq data to the combined HMBA BG/SEA-AD MTG atlas to assess changes in cellular proportions with respect to CPS. We utilized the scCODA model to evaluate the significance of changes in cellular abundance across AD pseudo-progression^50^. We found no significant changes in neurons across all mapped populations (Figure 3E), indicating that neither Aβ nor pTau pathology present in Ca is associated with neuronal loss in contrast to cortical regions^5,51,52^. However, we did observe changes in several non-neuronal populations with respect to CPS_AT8_. Similar to observations in MTG, we observed an increase in the protoplasmic astrocyte population Astro_2 with CPS_AT8_ (*mean p*_*inc*_ > 0.85; Figure 3E)^5,52^. We also found a Ca AT8-specific increase in the Micro-PVM_2_3-SEAAD population of microglia (Figure 3E). These microglia are not typically disease-associated, unlike microglia such as Micro-PVM_3-SEAAD^5,35,36,53^. We also observed a slight non-significant increase in Micro-PVM_2 (homeostatic microglia) across CPS in general. Changes in Astro_1, Oligo_2, and STR FS PTHLH-PVALB GABA shown in Figure 3E were not significant.

A significant proportion of differentially expressed genes (|*β*_*CPSx*_ | > 0.5, *p*_*adj*_ < 0.1) across cells are specifically in CPS_AT8_ and not in CPS_6E10_ suggesting that most gene changes are associated with pTau burden (Figure 3F). Surprisingly we find few differentially expressed genes (< 100 genes) across all neuronal cell types except for STR FS PTHLH-PVALB GABA neurons. Most differential expression was found in non-neuronal types such as microglia and oligodendrocytes. In spatial transcriptomics data, we observed highly variable white matter compartments across sampled regions in different donors (Supplemental Figure 1), suggesting that some of the differentially expressed genes in oligodendrocytes might in part be explained by such variation.

### Oligodendrocyte-rich regions spatially correlate with pTau and spatially disperse with A*β*

AD pathology is often localized to specific cell types or regions in the cortex, such as neurofibrillary tangles in pyramidal neurons^54^, or A*β* plaques spatially correlating with microglia^53,55^. While digital pathology can inform bulk associations with population and gene expression changes in snRNA-seq, such associations do not prove localization of AD associated pathology. To investigate whether the CPS_AT8_ associated abundance and gene expression changes could be explained by a localization of pTau, or if the lack of change of neuronal populations could explained by a lack of spatial correlation with pathology, we performed post-hoc immunostaining of Xenium spatial transcriptomic sections. Donors with high pathology load were defined as having at least 0.05% positive area for AT8 (4/10 spatial donors; 12/25 sections), or 6E10 (5/10 spatial donors; 16/25 sections). In these donors there was a broad correlation between both the machine segmented post-hoc immunofluorescence and the deep-learning segmented digital pathology (Supplemental Figure 2A, B). Neurofibrillary tangles are vanishingly rare in Ca, however, we do observe pTau aggregations resembling tangles in both D1 medium spiny neurons (MSN) and astrocytes (Supplemental Figure 2C, D), but the method did not allow for differentiating mature neurofibrillary tangles pathognomonic for AD from pre-tangles. Validating observations from digital pathology we found that a minority of AT8 signal was found in cell bodies as delineated by 18S RNA stain from the Xenium segmentation kit (Supplemental Figure 2E). We found that cell proximity to plaques and pTau differed from region to region (Supplemental Figure 2G, H) suggesting separate spatial compartments for AT8 and 6E10 pathological species.

To study the differential spatial distributions of the Ca we divided it based on oligodendrocyte density, as white and grey matter regions are interdigitated in basal ganglia and oligodendrocyte-dense regions tended to correspond to white matter (Figure 4A,B). We then compared white matter to regions with high MSN density which correspond to gray matter in the Ca (Figure 4E,F). As independent validation, we found that the regions with high oligodendrocyte density corresponded with white matter tracts as shown by the boundary stain (ATP1A1/E-cadherin/CD45) from the Xenium segmentation kit, which predominately labels non-neuronal membranes (Figure 4B). Comparing regions with a high density of oligodendrocytes (>10% coverage per bin) versus regions with oligodendrocyte sparsity we found an enrichment of pTau coverage in oligodendrocyte rich, white matter regions (Supplemental Figure 2F). Cellular proportions in distance bins reveal that oligodendrocytes represent 25% of all cells within 5 µm of pTau and dropping to 14% of all cells at 50 µm (Figure 4C). Neighborhood enrichment analysis based on pTau (*p* < 1 ∗ 10^−4^; Figure 4D) corroborated this finding with high enrichment (mean neighborhood enrichment score > 3) in all proximal spatial bins 0 – 40 *μm*. We note that pTau coverage is not exclusive to oligodendrocyte dense regions, and that a significant proportion of oligodendrocyte dense regions have little to no pTau in them (Supplemental Figure 2F; Figure 4B,F). Despite most differential expression changes being associated with CPS_AT8_, outside of oligodendrocytes no cell types have a specific association with pTau signal in the spatial transcriptomic data. Despite having neighborhood enrichments > 3, astrocytes and STR D1 and D2 MSNs associations with pTau vary greatly across donors. This can be in part explained by the sparseness of pTau inclusions in these cell types across donors (Supplemental Figure 2E) and the absolute number of MSNs in the spatial dataset. These patterns along with pTau found outside of cell bodies suggest that pTau is likely localized to myelinated tracts of either corticostriatal or pallidofugal axons in caudate.

**Figure 4.**
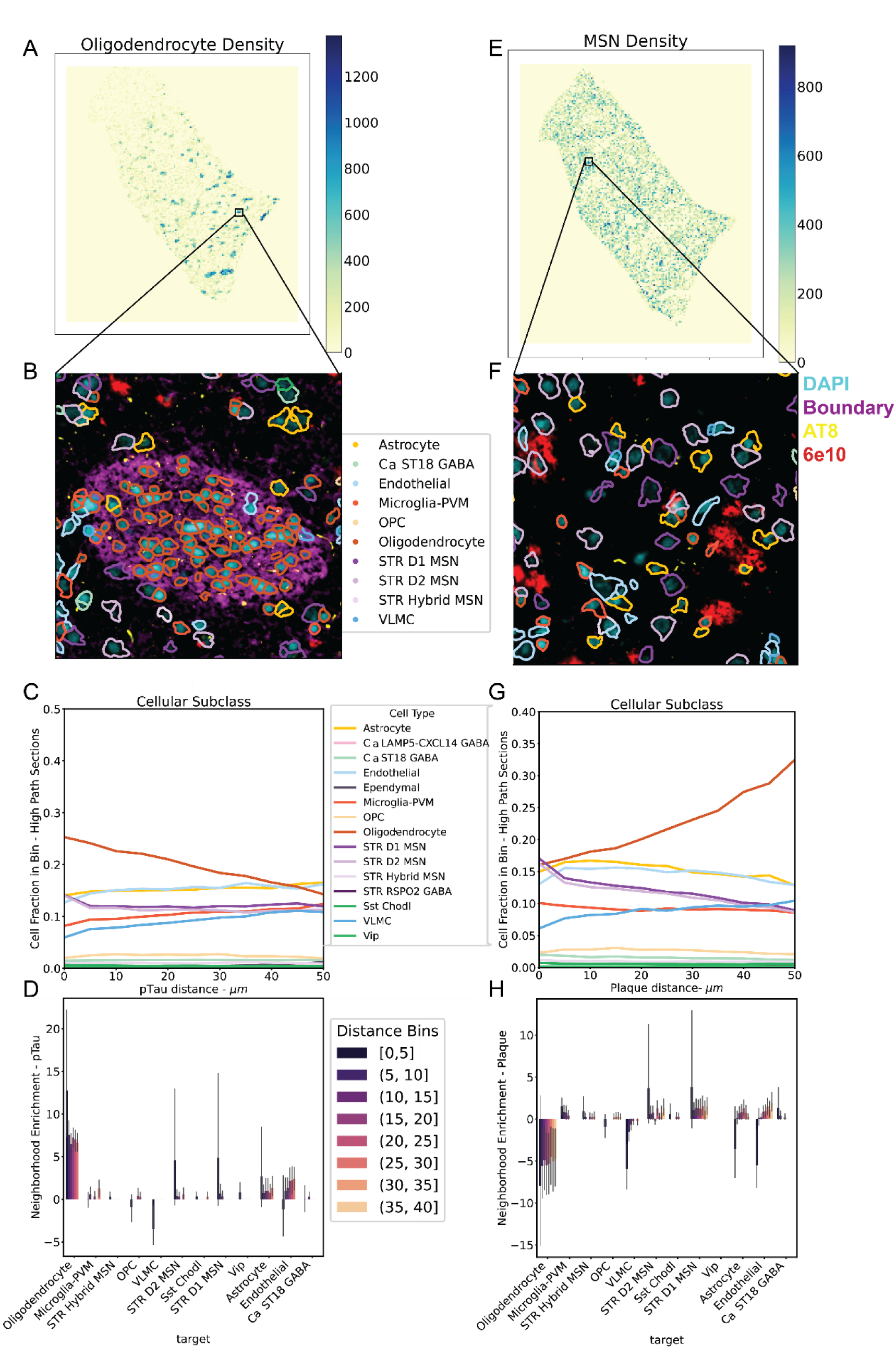
Oligodendrocytes and MSNs show opposing spatial relationships with respect to AD pathologies. (A-D) Spatial analysis of pTau and (E-H) Amyloid-*β*. 2D histogram of section colored by (A) oligodendrocyte and (E) MSN neuron density in each 2500 *μm*^2^bin from donor H21.33.002. (B,F) 62,500 *μm*^2^ inset of section showing spatial transcriptomic staining. AT8 (yellow) and 6E10 (red) post-hoc immunofluorescence. DAPI (cyan) and boundary stain (ATP1A1, E-Cadherin, CD45, magenta) from the Xenium segmentation kit. Polygons represent segmented cells. (C, G) Spatial distribution of proportions of cells within 5 *μm* bins pooled across high path donors. (D, H) Bar plots of neighborhood enrichment analysis scores. Error bars represent 95% confidence interval of scores across high pathology donors (high pTau load in D and high A*β* load in H). Bins values are in *μ*m.

In MSN rich regions, we found that Aβ plaques were the dominant pathology (Figure 4E, F). Looking at cellular proportions in binned distances from Aβ pathology, we find that oligodendrocytes represent 16% of cells within 5 µm of a plaque and increase to 32% of all cells at 50 µm. This trend is supported by neighborhood enrichment analysis showing significant spatial dispersion at 5-40 *μm*. (mean neighborhood enrichment score > 3). An opposing trend, we find that D1 and D2 MSNs represent 17% of cells within 5 *μm* of a plaque to 9% of all cells at 50 *μm* (Figure 4G). This result suggests that A*β* in the Ca is localized to the gray matter which is supported by previous in vivo measurements showing a spatial correlation between them in other cortical regions^56^.

Microglia and astrocytes are known to spatially associate with Aβ plaques during AD progression in cortex, where dense core and neuritic plaques are more prevalent^53,55^. Analysis of cellular proportions across distance bins and neighborhood enrichment reveals insignificant neighborhood enrichment (mean neighborhood enrichment score < 3) for microglia 0-5 *μm* and astrocytes from 20 – 25 *μ*m (Figure 4H). This suggests that diffuse plaques in Ca engender weak spatial associations with microglia and astrocytes.

### Ca microglia do not exhibit AD associated cortical microglia response to A*β* plaques

A lack of change in disease associated microglia (DAM), the increase in another microglial state (Micro-PVM_2_3-SEAAD), and the lack of broad microglial association suggest a distinct microglial response to AD pathology in the Ca. To assess this response, we first characterized Micro-PVM_2_3-SEAAD, and then performed in-depth spatial analysis of the A*β*-microglial niche.

To identify what pathways differentiate Micro-PVM_2_3-SEAAD from other microglia we performed gene set enrichment analysis on its marker genes using gene sets derived from factor analysis in Marshe et al (Figure 5A;Supplemental Figure 3A)^5,57^. The most highly enriched genes were components of the IL/IFN-*γ* signaling pathway with IL1B and FLT1 being the most highly expressed (Figure 5A). Notably, unlike cortical disease associated microglia (Micro-PVM_3-SEAAD), we observed negative enrichment of the NPY1R, Immuno-regulatory and APOE pathways. Given the role of the IL/IFN-γ pathway in promoting an inflammatory response and immune regulation, as well as NPY1R and APOE pathways in suppressing the same, this suggests that in the context of the neuropathology observed in caudate the Micro-PVM_2_3-SEAAD population may potentially be beginning a state-change toward disease-activated microglia^35,58–62^.

**Figure 5.**
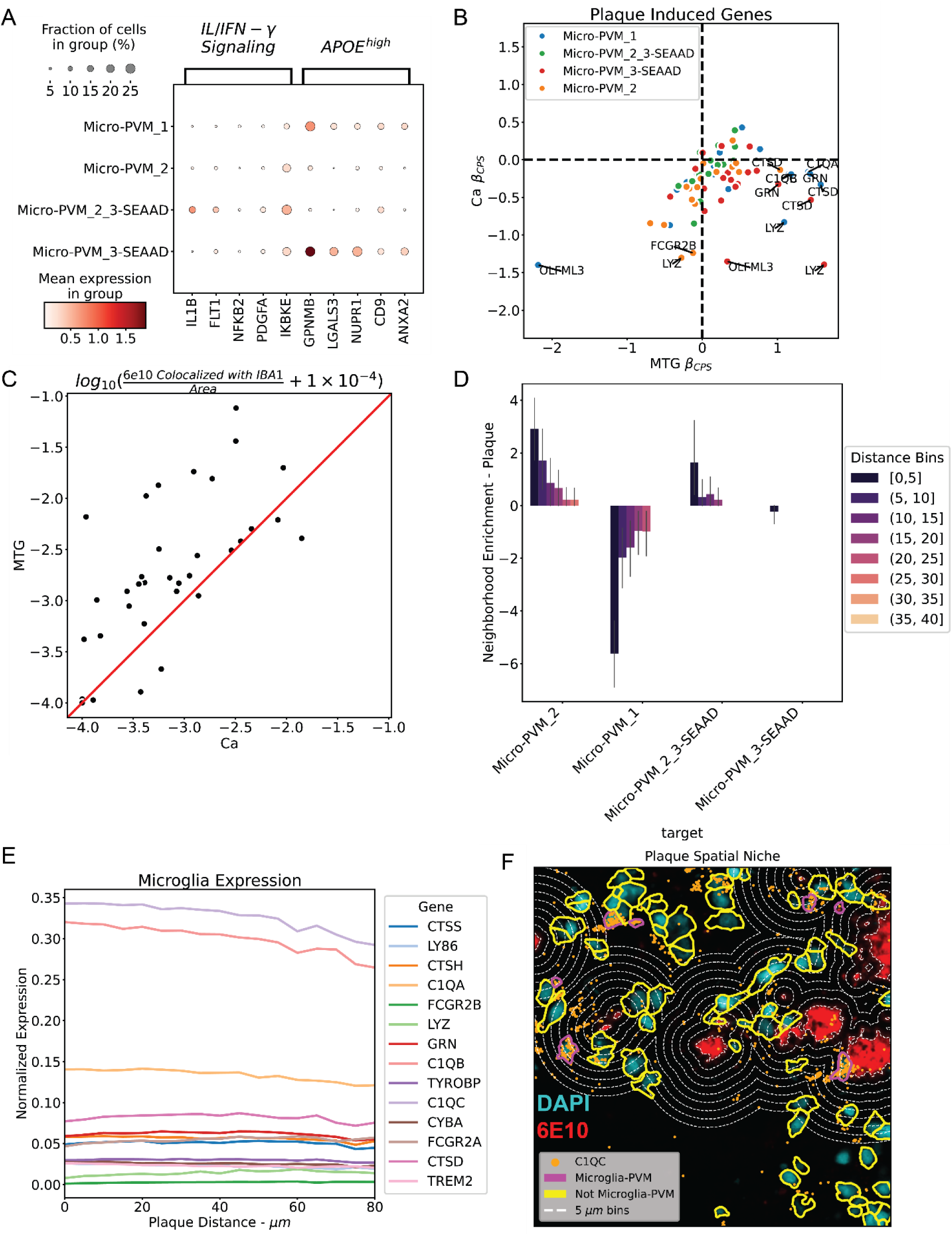
Microglia have weak response to, and spatial association with, Aβ plaques in Ca. (A) Dot plot where size is the proportion of microglia and saturation of red represents mean expression of microglia factor specific genes. (B) Scatterplot of plaque induced gene Nebula model *β*-coefficients in Ca vs. MTG from snRNA-seq. (*y* ∼ *Donor* + *Age* + *Sex* + *APOE*4 *Status* + 10*X Chemistry* + *CPS*). Dashed lines represent boundaries between positive and negative values. (C) Regression plot of 6E10 colocalized with IBA1 in Ca vs. MTG from digital pathology. Red line represents *y* = *x*.. (D) Bar plot of neighborhood enrichment scores with respect to A*β* plaques from spatial transcriptomics. Error bars represent 95% confidence interval across high A*β* load sections. (E) Line plot of mean plaque induced gene expression of 5 *μm* distance bins from A*β* plaques from spatial transcriptomics. Values pooled across all high A*β* load sections (F) Section inset (47350 *μm*^2^) from H20.33.002 showing spatial transcriptomic post-hoc immunofluorescence and expression of the plaque niche. DAPI is (cyan), 6E10 (red). *C1QC,* orange, was used as a marker of microglia. Segmentation polygons are colored either in magenta, microglia, or in yellow, non-microglia. Dashed contour lines represent distance bins for neighborhood analysis surrounding A*β* plaques.

To determine if the lack of the spatial association across all microglia were due to specific microglial subtypes associating with A*β*, we analyzed each subtype’s relationship to plaques and plaque-induced genes (PIGs)^53^. Using PIGs from the literature as a proxy for Aβ-induced response, we observed that genes in this set that were highly positively differentially expressed in MTG were negatively expressed in Ca (Figure 5B)^5,53^. Additionally, when looking at IBA1 duplexed with 6E10 in digital pathology, we observe IBA1 colocalization is more prevalent in MTG than in Ca for most donors (Figure 5C). We used spatial transcriptomics to study microglia in the plaque niche (Figure 5F) by measuring cellular proportions in distance bins (Supplemental Figure 3D) and performed neighborhood enrichment analysis (Figure 5D) on microglial subtypes. We find that there is association between homeostatic microglia (Micro-PVM_2) and Aβ (neighborhood enrichment score > 3) from 0-5 µm and an insignificant enrichment at 5-10 *μ*m (Figure 5D). We do not observe a significant enrichment of Microglia_2_3-SEAAD 0-5 *μ*m from A*β* plaques, and no PIGs were highly spatially enriched in microglia near Aβ plaques in Ca (Figure 5E).

Our findings point to a distinct and weaker response of microglia to Aβ plaques in Ca vs. other areas affected by AD. This may reflect the predominance of diffuse over compact and dense core plaques in Ca tissue. Additionally, this weaker response may in part explain the lack of changes in the neuronal populations of the Ca.

### IL1B+/FLT1+ microglia increase VEGFR1 signaling with AD

As *FLT1* is one of the prominent marker genes Micro-PVM_2_3-SEAAD, we looked into changes in genes relevant to the VEGFR1 signaling pathway. We observed increases in F*LT1*, *RAP1A*, *S1PR1* and a decrease in *GIPC1,* suggesting an overall increase in pathway activity (Figure 6A). All genes are specifically differentially expressed with respect to CPSAT8 and Micro-PVM_2_3-SEAAD except for FLT1, which is differentially expressed across all CPS values in both rather than Micro-PVM_2_3-SEAAD and Micro-PVM_2 (hoemostatic microglia; |*β*_*CPS*_| > 0.5; *p*_*adj*_ < 0.1; Figure 6A,Supplemental Table 4-6).

**Figure 6.**
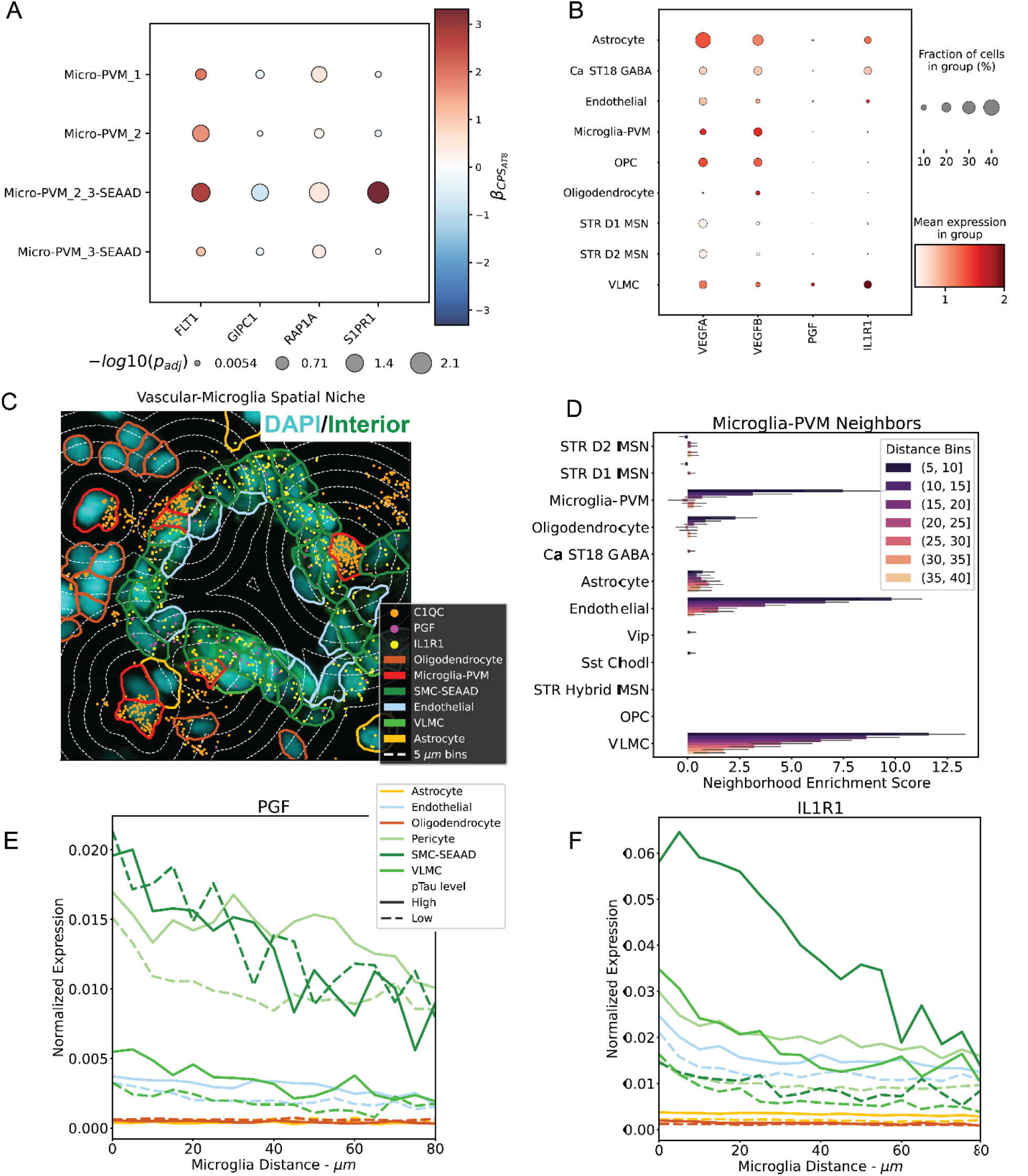
IL1B+/FLT1+ microglia increase VEGFR1 signaling with AD. (A) Dot plot of VEGFR1 signaling genes, where size is − log_10_( *p*_*adj*_), colored by *β*-coefficient for CPS_AT8_ from the Nebula model. Model equation: *y* ∼ *Donor* + *Age* + *Sex* + *APOE*4 *Status* + 10*X Chemistry* + *CPS*_*AT*8_. (B) Dot plot of snRNA-seq VEGFR1 signaling ligands genes, and the IL-1 receptor gene where size is proportion of cells colored by mean expression. (C) Section inset (11840 *μm*^2^) from donor H21.33.002 showing spatial transcriptomic segmentation stain and expression of the vascular – microglia niche. DAPI (cyan), Interior protein stain (alphaSMA/Vimentin, green). *PGF* (Magenta) and *IL1R1* (yellow) represented by dots. *C1QC (*orange) was used as a marker of microglia. Segmentation polygons are colored by cell type. Dashed contour lines represent distance bins for neighborhood analysis surrounding microglia. (D) Neighborhood enrichment scores for microglia. Error bars represent 95% confidence interval across all donors. Distance bin values are in *μ*m. (E, F) Lineplots of mean gene expression *PGF* and *IL1B* from spatial transcriptomics in 5 *μm* distance bins from microglia. Mean was calculated across either all high pTau load sections, or all low pTau load sections.

To determine which cell type may be acting upon upregulation of VEGFR1 pathway components, we looked at snRNA-seq measurements of canonical VEGFR1 ligand genes: *VEGFA*, *VEGFB*, and *PGF* (Figure 6B). We find that *VEGFA* is broadly expressed across cell types with the exception oligodendrocytes. *VEGFB* is also widely expressed. Finally, we find that *PGF* is highly expressed in a minority of VLMC cells. Having observed an increase in VEGF receptor expression in microglia and VEGF ligand expression across several other cell types, we wanted to validate potential ligand-receptor pairs in a spatial context (Figure 6C). To do that, we first performed neighborhood enrichment analysis to determine which cell-types tend to colocalize spatially with microglia (Figure 6D). While microglia tend to colocalize with themselves, they also strongly colocalize with vascular cell types (VLMC & endothelial cells). Further analysis revealed that *PGF* is enriched in proximity to Microglia in VLMCs, smooth muscle cells, and pericytes suggesting potential communication between the microglia and vascular cells through the VEGFR1 pathway (Figure 6E). In pericytes and VLMCs, both had elevated expression in donors with high pTau load.

As Microglia_2_3-SEAAD are enriched for *IL1B*, we also looked at the expression of its receptor, *IL1R1* in the snRNA-seq expression data and observe its enrichment in astrocytes, inhibitory interneurons, VLMCs, and a minority of endothelial cells (Figure 6B). We also observe that IL1R1 is enriched in proximity to microglia in all vascular cell types, suggesting another channel of communication through IL-1 signaling (Figure 6F). In all vascular cell types, we observe elevated IL1R1 expression in donors with high pTau burden. Taken together, these gene expression changes with AD progression, combined with a level of proximity conducive to active signaling suggest that microglia may be in communication with vascular cells in response to AD pathology in Ca.

### STR FS PTHLH-PVALB GABA exhibits global changes in key functional genes with respect to CPS_AT8_

When looking at the absolute numbers of significantly altered genes we find that fast-spiking PTHLH+ PVALB+ (FS PTHLH-PVALB) interneurons have the largest number of differentially expressed genes with respect to CPS_AT8_ (Figure 3F; Figure 7A). These are unique cells to the striatum that exhibit fast-spiking properties and have increased firing rates with respect to PVALB expression^64,65^. In the context of the basal ganglia, it has been shown in mice that related PVALB cells often receive excitatory inputs from the cortex and perform feed forward inhibition primarily on MSNs^66–68^.

**Figure 7.**
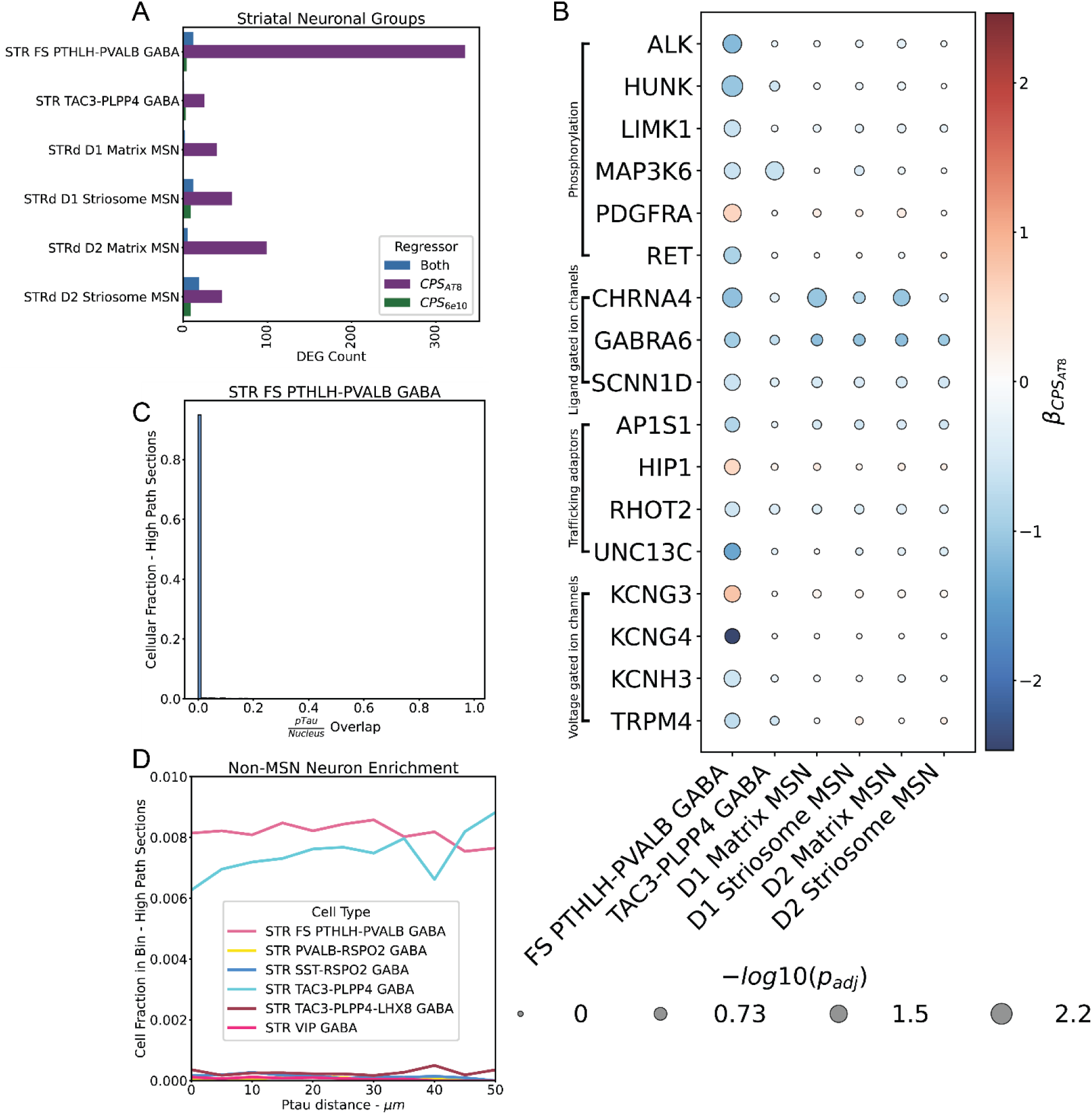
STR FS PTHLH-PVALB GABA exhibit global changes in key functional genes with respect to pTau load. (A) Bar plot number of significantly differentially expressed genes (DEG) across inhibitory neuronal types in Ca from snRNA-seq. (B) Dot plot differentially expressed genes grouped by gene sets, where size is − log_10_( *p*_*adj*_) and color is *β*-coefficient for CPS_AT8_ from Nebula model (*y* ∼ *Donor* + *Age* + *Sex* + *APOE*4 *Status* + *method* + *CPS*_*AT*8_). (C) Histogram of overlap distribution between STR FS PTHLH-PVALB GABA nuclei and pTau from spatial transcriptomics. (D) Line plot of cellular proportion of 5 *μm* distance bins from pTau from spatial transcriptomics. Values pooled across all high pTau load sections.

Downregulation (*β*_*CPSAT*8_ < −0.5; *p*_*adj*_ < 0.1) of genes in several pathways suggests altered functional activity in FS PTHLH-PVALB (Figure 7B). We found genes differentially expressed specifically in FS PTHLH-PVALB encoding voltage-gated and ligand-gated ion channels (*KCNH3*, *KCNG4,* and *TRPM4)* were down regulated. Additionally, we found that genes associated with trafficking adaptors (*AP1S1*, *RHOT2*, and *UNC13C)* were downregulated. While not specific to FS PTHLH-PVALB (this effect was observed in D1 and D2 matrix MSNs) we find that the ligand-gated ion channel gene *CHRNA4* is down regulated with respect to CPS_AT8_. This is similar to changes in CHRNA4 in MTG inhibitory neuronal types^5^. We found exceptions to the general trend of down regulation (*β*_*CPS*_ > 0.5; *p*_*adj*_ < 0.1) in these genes, namely upregulation of *HIP1* trafficking adaptor gene and the voltage gated ion channel gene *KCNG3*.

To see if the effect was spatially localized or global with respect to pTau we looked for any associations with pTau in the high pTau load Xenium sections (Figure 7C,D). We find that the vast majority of FS PTHLH-PVALB interneuron cell bodies do not contain any pTau in them (Figure 7C). Additionally, we find no spatial association specific to FS PTHLH-PVALB interneurons (Figure 7D). We find little evidence that these gene expression changes are a direct result of proximity to local pathology.

## DISCUSSION

We performed snRNA-seq and spatial profiling of caudate head tissue in a select group of AD donors to determine which cell types might be affected or changing with disease progression, what neuropathological components are present and any changes in gene expression that accompany AD in this region. Neuropathology present in the caudate nucleus differs from that in the cortex of the same donor in that while affected cortical regions contain a mixture of dense-core, compact and diffuse plaques, caudate nucleus contains primarily diffuse plaques. In addition, pTau is present but predominately in processes in neuropil thread form and not perinuclear tangle form as in affected cortical regions. Unlike our previous observations in cortex^5^, caudate neuronal proportions are unchanged even in donors with high pathological burden. However, non-neuronal populations do change in relationship to pathology in caudate. Specifically, we observe differential gene expression and abundance changes in specific microglial populations indicating an immune response unlike that of a classical disease associated microglia. Spatial transcriptomics results reveal that microglia most closely associate with each other and vascular cells. Taken with our finding that components of interleukin and VEGF signaling are upregulated in both microglia and vascular cells with pathological progression, there may be AD-related changes occurring in the vasculature. However, we do not observe the classical arrangement of microglia, astrocytes, and oligodendrocytes proximal to plaques in Ca as has been observed in cortical regions^53^. Alternatively, microglial activation and association with vessels in Ca may reflect a reaction to vascular pathology not directly related to AD, such as arteriolosclerosis. Some Ca neuronal populations undergo significant differential gene expression in response to pTau pathology despite the dearth of intracellular tangles. Notably, fast-spiking interneurons exhibit alterations in receptor tyrosine and serine threonine kinases, ion channels and synaptic protein expression that likely impact function. Taken together, these results suggest that the nature of the AD pathology present in caudate combined with the primarily GABAergic neuronal circuitry produce unique cellular and circuit responses.

### Neurofibrillary tangles and neuritic plaques are rare in the Ca

We observe that Ca exhibits a distinct pathological make up in AD. While there is a correlation in overall 6E10 and AT8 signal for Ca versus MTG, there are 1-2 fold fewer dense-core plaques and neurofibrillary tangles in Ca than in MTG. We confirm previous findings that Aβ plaques were predominantly diffuse in Ca^39–41^. Surprisingly, we find that pTau in Ca exists mostly in the form of neuropil threads outside of cell bodies and enriched in white matter tracts. Thus, we note that Ca gives us a unique look at the effect of these specific pathological species in late-stage donors independent of the effects of other hallmarks of AD pathology.

We speculate that the observed Ca-specific AD pathology has a weaker pathogenic profile. Diffuse plaques in the cortex are marked by their lack of glial response and are not associated as strongly with dementia as focal plaques^43,69^. Additionally, diffuse plaques are often observed in pre-clinical aging brains and are hypothesized to develop into dense core and then neuritic plaques^70,71^. This lack of neuritic plaques is refltected in the lack of cellular pTau and the enrichment of pTau and A*β* plaque being enriched in different spatial compartments in the Ca. Additionally, neuropil threads are often associated with earlier, non–clinical stages of AD pathology^72–76^. We conjecture that these associations with earlier or non-clinical forms of AD are reflected in the lack of changes in neuronal and oligodendrocyte abundances, and the distinct response of microglia in the Ca.

### Diffuse amyloid pathology in caudate is weakly associated with glial response

Glial response to Aβ in Ca was muted when compared to studies in other AD-affected regions. In snRNA-seq we observe little change in key reactivity genes in either astrocytes (*GFAP, CD44, CHI3L1)* or microglia (*FCGR2A*, *C1QA-C*) (Supplemental Table 4-6)^5,53,77^. Additionally, we do not find a change of *SERPINH1* in the protoplasmic astrocytes as previously observed (Supplemental Table 4-6)^22^. In digital pathology and snRNA-seq we find weaker signal for colocalization of microglia with A*β* plaques than in MTG. However, we observe spatial enrichment of homeostatic microglia with respect to Ca Aβ plaques suggesting that some chemotaxis might be occurring. Microglia near Aβ plaques do not exhibit substantially different expression of plaque induced genes^53^. We surmise that diffuse Aβ plaques in the absence of dystrophic neurites are not enough to elicit the glial response associated with standard AD progression. This muted response may, in part, explain the lack of neuronal cell death.

### FLT1+/IL1B+ microglia and protoplasmic astrocytes are associated with global pTau burden

All abundance changes and the most significant expression changes were associated with CPS_AT8_. We observe significant abundance increases in protoplasmic astrocytes and microglia enriched in IL/VEGF signaling. Additionally, we observe marked changes in VEGF signaling expression in microglia and changes in key ion channel genes in striatal fast-spiking PTHLH+/PVALB+ GABAergic neurons. However, we observe neither an enrichment of neurofibrillary tangles nor a spatial enrichment near neuropil threads of impacted cell types (Figure 3D). While neurofibrillary tangles in D1 MSNs were observed in the tissue, these were rare occurrences even in high Braak stage donors. This lack of tangles may reflect specific resistance to pTau accumulation in the Ca. In the striatal fast-spiking neurons, we find a transcriptional decrease for receptor tyrosine and serine threonine kinases (e.g. HUNK, RET, LIMK1, and ALK) (Figure 7B). However, these changes were not universal across other neuron types. Additionally, we did not observe significant changes in the expression of genes usually associated with pTau accumulation such as *CUL5* and *MAPK* (Supplemental Table 5)^78,79^. Alternatively, assuming pTau spreads through neuronal connections as hypothesized widely and given the observation that pTau is enriched in white matter, pTau spread to the Ca is so late in AD progression that donors die before the region can be seeded^80–82^. The lack of spatial enrichment of neuropil threads to impacted cell types suggest that the effect of pTau on the Ca is global rather than localized to pathology.

### Microglia increase VEGFR1 signaling with AD

We found a robust increase in VEGFR1/FLT1 with respect to pTau burden in Ca specific disease associated microglia. Several studies in the ROS/MAP cohort have associated VEGFR1 signaling genes with cognitive decline, A*β* and/or pTau burden^83–85^. Studies using *in vitro* microglia and *in vivo* rat brains have implicated A*β*_1−42_, one of the primary components of A*β* plaques, in the increased expression of FLT1 and that FLT1 in turn was important in the recruitment of microglia to A*β* plaques^86,87^. We observe a significant change (*p*_*adj*_ < 0.1, *β*_*CPS*_ > 0.5) in homeostatic microglia for CPS_6E10_ along with A*β* plaque neighborhood enrichment (mean neighborhood enrichment score > 3), corroborating this. Interestingly, a recent publication found that expression of other VEGF signaling genes (*FLT4 and NRP2*) were negatively correlated with Aβ and pTau accumulation in Ca using bulk sequencing^88^. However, in snRNA-seq, we find no differential expression of these genes in microglia specifically (Supplemental Table 4-6). Increases in VEGF signaling prompted us to investigate a potential relationship between microglia and vascular cells. It has been established that microglia and micro-vasculature in the brain physically interact^89,90^. Additionally, it has been shown that through the action of chemokines like FLT1 and IL1B microglia can induce angiogenesis^91^. Utilizing spatial transcriptomic analysis, we recapitulated colocalization of microglia and vasculature. Additionally, we found that *FLT1* ligand, *PGF*, and *IL1B* receptor, *IL1R1*, are both enriched in vasculature proximal to microglia in high path donors. This provides strong evidence that VEGF and IL signaling is enriched across AD, implying altered behavior in vascular associated microglia. Microvascular alterations, including vascular remodeling are a well-known phenomenon in AD^92^, and vascular dysregulation may be among the early changes in AD pathogenesis^93^. Future characterization of this niche in other regions may reveal an underlying mechanism relating vascular disease to AD, as vascular disease is a common comorbidity of AD.

### Localization of pTau in white matter tracts suggests potential circuit dysfunction reflected in STR FS PVALB-PTHLH GABA gene expression changes

Interpretation of pTau enrichment in oligodendrocyte dense regions is complicated by the circuits of the basal ganglia and differential sampling of white matter tracts. We find variable counts of oligodendrocyte dense regions from donor to donor (Supplemental Figure 1), suggesting potentially differential donor sampling of fasciculate axons even when sampling closely spatially apposed sections of different donors. While we do observe differential expression in oligodendrocytes, it is unclear if this expression is a reflection of true differential expression in AD, or differential sampling of oligodendrocyte compartments in the Ca. Additionally, given that pTau is mostly localized to the neuropil, and that there is an enrichment in oligodendrocyte dense regions corresponding to white matter tracts, much of the signal we are observing could be originating from projecting neurons from different regions of the cortex or the basal ganglia (Figure 1A)^94^. In the context of AD, cortico-striatal connections are the strongest candidate, given that the dorsal prefrontal cortex of the brain would be experiencing high pTau load in the most affected donors^95^. However, pTau has also been observed in basal ganglia areas projecting intraregionally in the context of other diseases such as Parkinson’s and progressive supranuclear palsy^96,97^. However, discerning the origins of fasciculated axons such as cortico-striatal, striato-pallidal or nigro-striatal in the caudate nucleus is beyond the scope of this paper and would likely need validation through non-human primate modeling of AD.

Basal ganglia circuitry may also explain the changes in expression of functionally important genes in FS PTHLH-PVALB interneurons. There are known cortico-striatal connections between projecting IT neurons of the cortex and striatal fast-spiking neurons^66^, and previous studies show that such excitatory neurons are subject to neurodegeneration^5,98^. Our past work and that of others has uncovered unique vulnerabilities in multiple inhibitory interneuron types in neurodegenerative and other psychiatric diseases^5,99^. Furthermore, PVALB-expressing interneurons are exquisitely sensitive to changes or reductions in network activity because they are often fast-spiking, metabolically demanding cells whose molecular identity and firing properties are highly activity-dependent^100–102^. PVALB expression, firing rate and synaptic inputs all adapt to the level of excitatory drive, so chronic activity loss causes downregulation of PVALB-related proteins, reduced fast-spiking output and impaired inhibition – changes that serve as a readout for and a contributor to circuit dysfunction in neurodegenerative disease. On a more granular level, these vulnerabilities stem from high energy/ion-handling requirements, precisely balanced expression of sodium and calcium channel complements and sensitivity to oxidative stress and synapse loss^64,100^. In fact, PVALB-centric studies of mouse AD models have shown that PVALB cell dysfunction is an early event contributing to cognitive decline and that rescuing PVALB activity can restore network rhythms and improve memory^103–105^. Here we observe that functional genes associated with ion channels and trafficking adaptors were inhibited with AD progression in caudate PVALB neurons, suggesting that these cells may have altered intrinsic physiological properties. Thus, we speculate that altered upstream signaling from cortical neurons projecting to FS PTHLH-PVALB interneurons may cause dysfunctional neuronal firing in said interneurons. Taken together, differential gene expression within and disease-affected input to PTHLH fast spikers may portend a shift in inputs to MSNs in the Ca in this context^106^.

### Towards understanding Ca function in AD and other neurodegenerative diseases

Our paper represents a descriptive, cellular resolution look at the impact of AD pathology in the Ca and is thus a powerful resource for future neurodegenerative studies. In AD, this dataset is a useful comparator due to its unique pathological make up of mostly diffuse Aβ and neuropil threads. Ca is a useful contrast to other regions of the brain such as the hippocampus and the cortex that exhibit more mature forms of pathology such as neuritic plaques and neurofibrillary tangles. Additionally, due to its downstream connections with other AD affected regions, such as dorsal prefrontal cortex, this work reveals how AD pathology might spread to the dorsal striatum in high Braak and Thal donors. The Ca is also impacted by other neurodegenerative diseases comorbid with AD such as Lewy Body Dementia and Huntington’s disease^30–33^. How the unique pathological proteins present in each of these diseases contribute to brain dysfunction individually remains an open question. Our cohort specifically aims to minimize these comorbidities, and thus focus on effects from AD and aging. Thus, our dataset represents an AD-specific diseased control in the Ca, and is uniquely poised to help distinguish the unique and synergistic effects of neurodegenerative diseases of the dorsal striatum.

## RESOURCE AVAILABILITY

### Lead contact

- Requests for further information and resources should be directed to and will be fulfilled by the lead contact, Jennie Close.

### Materials availability

- This study did not generate new unique reagents. Data and code availability
- FASTQ files containing the sequencing data from the snRNA-seq, snATAC–seq and snMultiome assays are available through controlled access at Sage Bionetworks (accession no. syn26223298). Instructions for access to data on the AD Knowledge Portal is provided by Sage Bionetworks.
- Anndata h5ad file containing nuclei-by-gene matrices with counts for snRNA-seq and snMultiome data deposited on the [Open Data Registry](https://registry.opendata.aws/allen-sea-ad-atlas/) in an AWS bucket (sea-ad-single-cell-profiling) as AnnData objects (h5ad files).
- Anndata h5ad file containing cell-by-gene matrices with counts for 10X Xenium spatial transcriptomics is available on the [Open Data Registry](https://registry.opendata.aws/allen-sea-ad-atlas/) in an AWS bucket (sea-ad-spatial-transcriptomics).
- Spatialdata zarr files containing images, labels, polygons, and cell-by-gene matrices for each section of the 10X Xenium spatial transcrptomics data is available on [Open Data Registry](https://registry.opendata.aws/allen-sea-ad-atlas/) in an AWS bucket (sea-ad-spatial-transcriptomics).
- 10X Xenium spatial transcriptomic output files, with post-hoc immunostaining images and alignments are available on the [Open Data registry](https://registry.opendata.aws/allen-sea-ad-atlas/) in an AWS bucket (sea-ad-spatial-transcriptomics).
- All original code pertaining to the reproduction of figures has been deposited at: https://github.com/AllenInstitute/2026_SEA_AD_Ca_Atlas_Paper_Figure_Reproducibility
- All original code pertaining to spatial analysis has been deposited at: https://github.com/AllenInstitute/SEA_AD_CaH_spatial_transcriptomics
- All original code pertaining to the interative scANVI implementation has been deposited at: https://github.com/AllenInstitute/SEA_AD_CaH_iterative_scANVI
- All original code pertaining to the differnential abundance implementation in pertpy has been deposited at: https://github.com/AllenInstitute/SEA_AD_CaH_perpty/
- All code pertaining to the differential expression implementation in Nebula has been deposited at: https://github.com/AllenInstitute/SEA_AD_CaH_Nebula/
- Any additional information required to reanalyze the data reported in this paper is available from the lead contact upon request.

## ACKNOWLEDGMENTS

The SEA-AD consortium is supported by a National Institute on Aging (NIA) grant no. U19AG060909. The study data were generated from postmortem brain tissue donated to the University of Washington BRaIN laboratory and Precision Neuropathology Core, which is supported by the UW ADRC (NIA grant no. P30AG066509, previously no. P50AG005136), the ACT study (NIA grant no. U19AG066567) and U24AG072458, U24NS135561, U24NS133945, U24NS133949, RF1AG065406, R01NS105984, R01AG60942 and UM1MH130981. Additionally, ACT data collection for this work was supported, in part, by prior funding from the NIA (no. U01AG006781) and the Nancy and Buster Alvord Endowment (to C.D.K.). All statements in this manuscript, including its findings and conclusions, are solely those of the authors and do not necessarily represent the views of the National Institute on Aging or the National Institutes of Health. We thank the participants of the ADRC and the ACT study for the data they have provided, and the many ADRC and ACT investigators and staff who steward that data. You can learn more about the UW ADRC at https://depts.washington.edu/mbwc/adrc and ACT at https://actagingstudy.org/. https://portal.brain-map.org/explore/seattle-alzheimers-disease

## AUTHOR CONTRIBUTIONS

O.Z.K.,K.J.T., E.S.K., M.I.G., N.P.,B.L.,C.D.K., E.S.L.,J.L.C. Conceptualization; R.D.H.,J.T.M., E.C.G., M.X., T.S.B., V.M.R., E.J.M., S.L.O., H.H., N.P., C.D.K., D.A.M., N.M., N.V.C., P.O., J.N.,J.C., J.G., A.B., R.C., M.T., T.C., A.T., D. B., J.G.,R.F.,K.S.,N.D., S.B., A.O., A.R, C.A.R., A.Ay., M.B., A.H.,J.A., N.M.G., J.W., P.C.K., T.G., Data Curation, O.Z.K., K.J.T., B.L., A.Ag., N.P., M.X., M.I.G., J.L.C, formal analysis, C.D.K., E.S.L., J.L.C., Funding Acquisition. O.Z.K., K.J.T., B.L., A.Ag., G.E.M., J.H., M.I.G., J.L.,J.G., Software; Resources; E.S.K. project administration; O.Z.K., J.L.C.,, M.H. Visualization; O.Z.K., J.L.C. writing – original draft; B.L., K.J.T., E.S.K., M.I.G., N.P., R.D.H., J.A.M., M.H., C.D.K., E.S.L., writing – review & editing

## DECLARATION OF GENERATIVE AI AND AI-ASSISTED TECHNOLOGIES

During the preparation of this work, the authors utilized Microsoft Copilot to help generate code for analysis. All subsequent code was reviewed and tested by the authors. The authors full responsibility for the content of the publication.

## SUPPLEMENTAL INFORMATION

**Document S1** Contains Supplemental Figure 1 and 2.

**Table S1** List of genes included in the gene panel of the xenium experiments along with the number of probes for each gene.

**Table S2** Digital pathology measurements for AT8, 6E10, and TDP-43 in caudate and middle temporal gyrus.

**Table S3** Differential abundance analysis results from scCODA.

**Table S4-6** Differential expression analysis results from NEBULA, across global, AT8 and 6E10 CPS respectively.

**Table S7** CPS measurements along with pathology measurements used to infer CPS for each donor.

**Table S8** Donor cohort demographic and neuropathological metadata.

## STAR★METHODS

### KEY RESOURCES TABLE

The items in the key resources table (KRT) must also be reported alongside the description of their use in the method details section. Literature cited within the KRT must be included in the references list. Please **do not edit the headings or add custom headings or subheadings** to the KRT. We highly recommend using [RRIDs](https://scicrunch.org/resources) as the identifier for antibodies and model organisms in the KRT. To create the KRT, please use the template below or the [KRT webform](https://star-methods.com/). See the more detailed [Word table template](http://www.cell.com/pb-assets/journals/research/cell/methods/table-template1.docx) document for examples of how to list items.

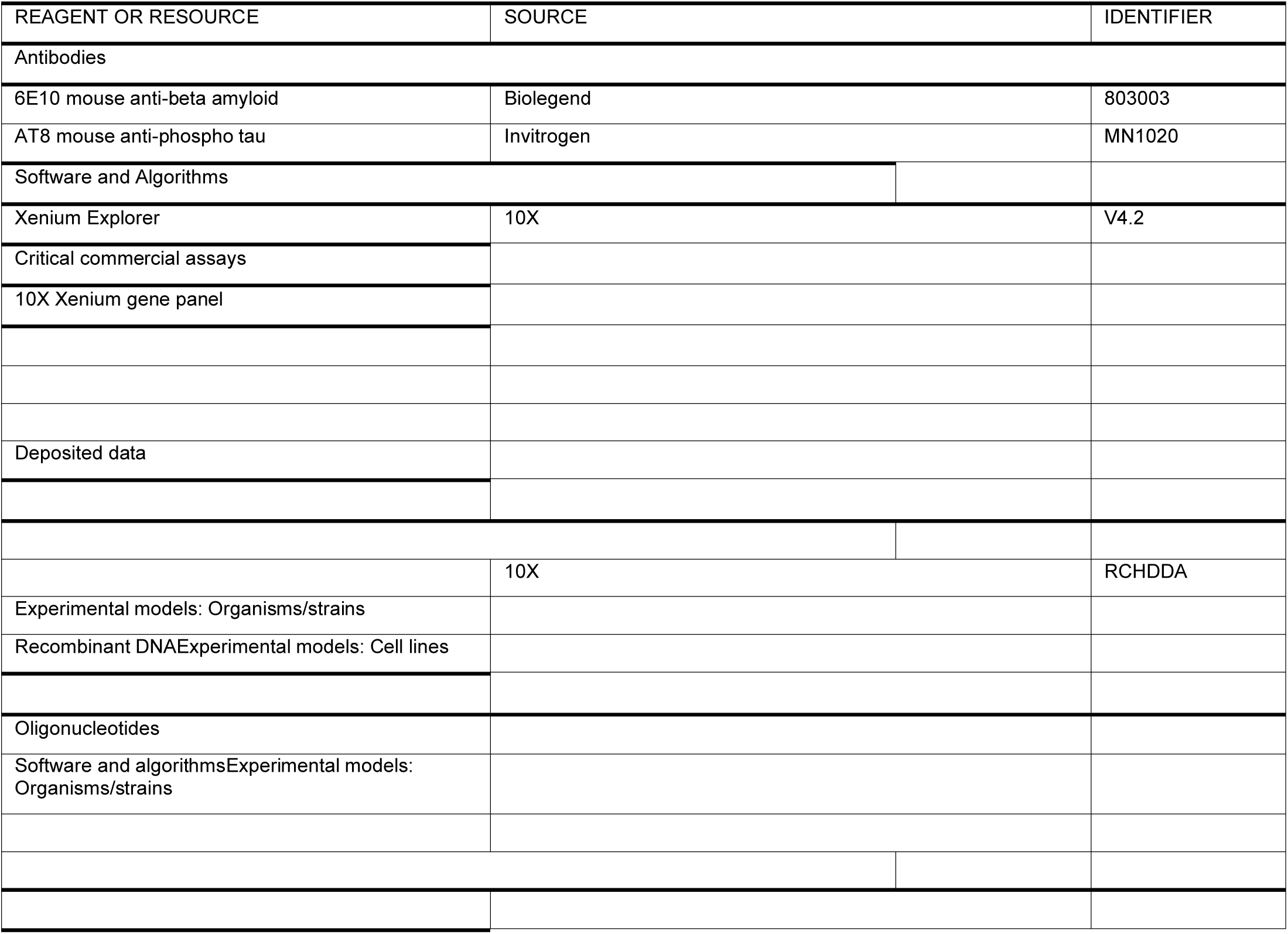
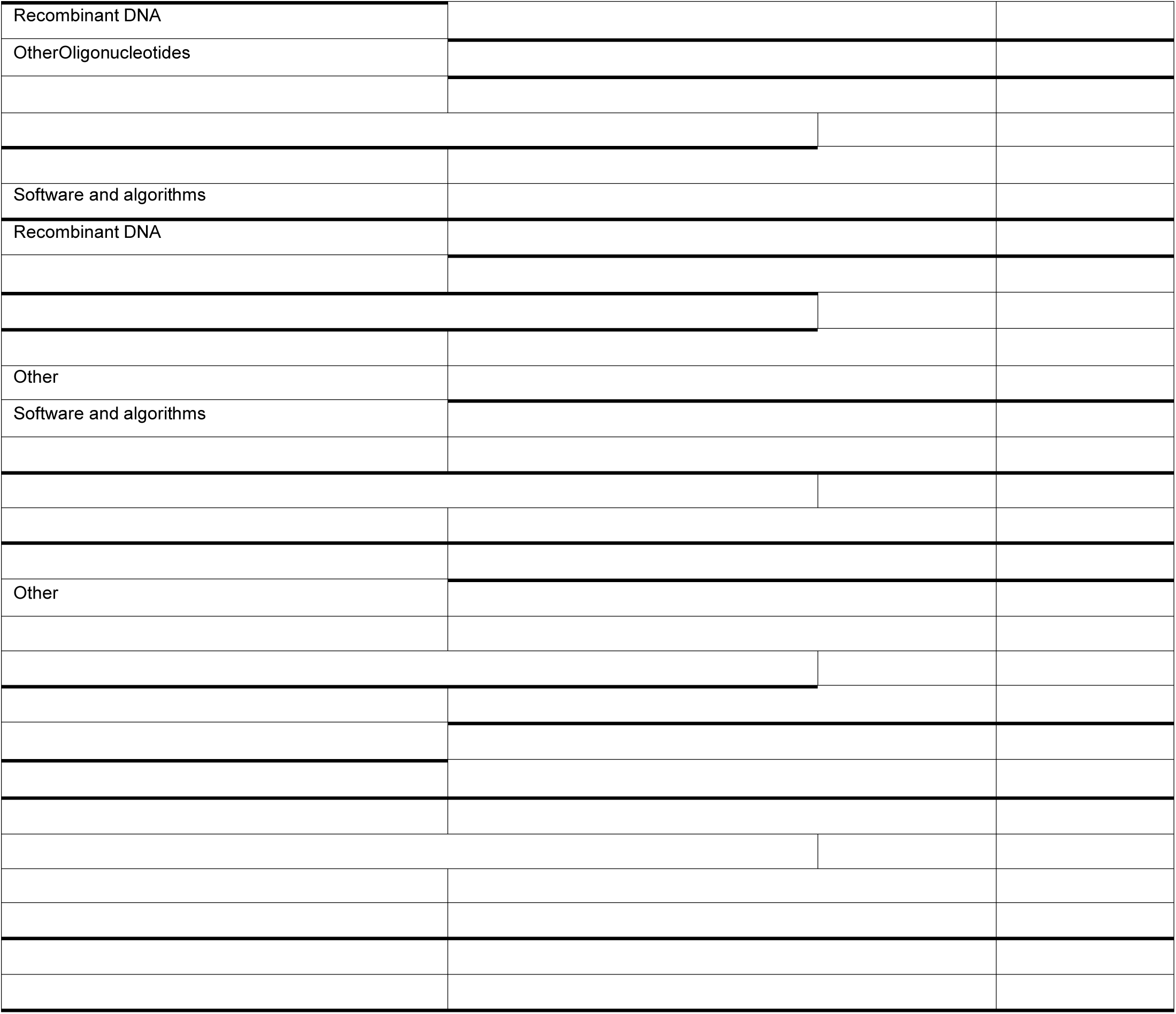

### METHOD DETAILS

#### SEA-AD cohort selection and brain tissue collection

Brain specimens were donated for research to the University of Washington (UW) BioRepository and Integrated Neuropathology (BRaIN) laboratory from participants in the Adult Changes in Thought (ACT) Study and the University of Washington Alzheimer’s Disease Research Center (ADRC). The study cohort was selected based solely on donor brains undergoing precision rapid procedure (optimized tissue collection, slicing, and freezing) during an inclusion time period at the start of the SEA-AD study, as previously described^5^. Donors with a diagnosis of frontotemporal lobar degeneration (FTLD), Down’s syndrome, amyotrophic lateral sclerosis (ALS) or other confounding degenerative disorder (not including Lewy body disease, limbic-predominant TDP-43 encephalopathy, or microvascular brain injury) were excluded. The cohort was chosen in this manner to represent the full spectrum of Alzheimer’s disease neuropathology, with or without common comorbid age-related pathologies.

The ACT study is a community cohort study of older adults from Kaiser Permanente Washington (KPW), formerly Group Health, in partnership with the UW. The ACT study seeks to understand the various conditions and life-long medical history that can contribute to neurodegeneration and dementia and has been continuously running since 1994, making it the longest running study of its kind. In 2004, ACT began continuous enrollment with the same methods to replace attrition from dementia, dropout, and death, ensuring a consistent cohort of ≥2,000 at risk for dementia. Total enrollment is nearing 6,000, with over 1,000 incident dementia cases; more than 900 have had autopsies to date with an average rate of approximately 45-55 per year. The study completeness of the follow up index is between 95 to 97%. Subjects aged 65 or older without dementia are invited to enroll by random selection from the greater Seattle area patient population of KPW Seattle and undergo bi-annual study visits for physical and mental examinations. In addition to this study data, the full medical record is available for research through KPW. Approximately 25% of ACT autopsies are from people with no MCI or dementia at their last evaluation; roughly 30% meet criteria for MCI, and roughly 45% meet criteria for dementia. Thus, the ACT study provides an outstanding cohort of well-characterized subjects with a range of mixed pathologies including many controls appropriate for this study. Approximately 30% of the ACT cohort consents to research brain donation upon death, and tissue collection is coordinated by the UW BRaIN lab, which preserves brain tissue for fixed, frozen, and fresh preparations (described below), as well as performing a full post-mortem neuropathological examination and diagnosis by Board-certified neuropathologists using the NIA-AA and other relevant, current guidelines.

The UW Alzheimer’s Disease Research Center (ADRC) has been continuously funded by NIH since 1984. It is part of a nationwide network of ADRCs funded through the NIA and contributes uniquely to this premier program through its vision of precision medicine for AD: comprehensive investigation of an individual’s risk, surveillance with accurate and early detection of pathophysiologic processes while still preclinical, and interventions tailored to an individual’s molecular drivers of disease. Participants enrolled in the UW ADRC Clinical Core undergo annual study visits, including mental and physical exams, donations of biospecimens including blood and serum, and family interviews. The UW ADRC is advancing understanding of clinical and mechanistic heterogeneity of Alzheimer’s disease, developing pre-clinical biomarkers, and, in close collaboration with the ACT study, contributing to the state of the art in neuropathological description of the disease. For participants who consent to brain donation, tissue is also collected by the UW BRaIN lab, and is preserved and treated with the same full post-mortem diagnosis and neuropathological work up as described above.

Human brain tissue was collected at rapid autopsy (postmortem interval <12 hours, mean close to 7 hours, **Supplemental Figure. 1a**). One hemisphere (randomly selected) was embedded in alginate for uniform coronal slicing (4mm), with alternating slabs fixed in 10% neutral buffered formalin or frozen in a dry ice isopentane slurry on Teflon-coated plates.

##### Single and duplex immunohistochemistry (IHC) for quantitative neuropathology

Tissue for quantitative neuropathology from each region were sampled from fixed slabs embedded in paraffin. The tissue blocks were sectioned (cut at 5 µm), deparaffinized by immersion in xylene for 3 minutes, 3 times. Then, rehydrated in graded ethanol (100%, 3x, 96%, 70% and 50% for 3 minutes each) and washed with TBST (Tris Buffered Saline with 0.25% Tween) twice for 3 minutes. The slides were immersed in Diva Decloaker 1x solution (Biocare Medical, DV2004) for heat-induced epitope retrieval (HIER) using the Decloaking Chamber at 110C for 15 minutes for most of the antibodies. For the alpha-Synuclein protein detection, enzymatic antigen retrieval with protein kinase is used. After the HIER is completed, the slides are cooled for 20 minutes at RT. Afterward, the slides are washed with TBST for 5 minutes, twice.

Chromogenic staining was performed using the fully automated BioCare Medical intelliPATH®. Blocking with 3% hydrogen peroxide, Bloxall (Vector Labs), Background punisher (BioCare Medical), and levamisole (Vector labs) is performed to avoid any cross-reactivity and background. The following primary antibodies are used for the first target protein at the dilutions indicated: NeuN (1:500, A60, Mouse, Millipore MAB5374), pTDP-43 (1:1000, Ser409/Ser410, ID3, Rat, Biolegend, 829901), Aβ (1:1000, 6E10, Mouse, Biolegend 80303), Alpha-Synuclein (1:200, LB509, Mouse, Invitrogen 180215) and GFAP (1:1000, Rabbit, DAKO, Z033401-2). Following primary antibody incubation sections were washed 4×2 minutes with TBST and stained with species-appropriate secondary antibody conjugated to a Horseradish Peroxidase (HRP, MACH3-Mouse (M3530), and MACH-Rabbit (M3R531), BioCare Medical). Sections were washed 2×2 minutes with TBST and the antibody complex is then visualized by HRP-mediated oxidation of 3,3’-diaminobenzidine (DAB) by HRP (brown precipitate). Counterstaining is done with hematoxylin after the DAB reaction.

In the case of a duplex IHC (Aβ and pTDP43), the slides were washed 18×2 minutes in TBST and then incubated with primary antibodies at the dilutions indicated after the DAB reaction: Iba1 (1:1000, Rabbit, Wako, 019-19741) and pTau (1:1000, AT8, Mouse, Thermofisher, MN1020), washed as above and stained with species-appropriate secondary antibodies conjugated to an Alkaline Phosphatase (AP, MACH3-Mouse (M3R532) MACH3-Rabbit (M3R533), Biocare Medical). The complex was then visualized with the intelliPATH® Ferangi Blue reaction kit (IPK5027, Biocare Medical) (blue precipitate). Once staining is completed, the slides were removed from the automated stainer and immersed in TBST, 3 minutes, then dehydrated in graded ethanol (70%, 96%, 100%, 2x) for 3 minutes and xylene (or xylene substitute in the case of double IHC), 3 times each for 3 minutes. Finally, coverslipping is carried out with a Tissue-Tek automated cover slipper (Sakura).

#### Acquisition and quantification of whole slide neuropathology images

Whole slides stained with IHC were scanned on the Aperio AT2 digital scanner (Leica), which captures sequential images of a 20x field of view, using slide settings optimized for our IHC protocols which are subsequently assembled or stitched into whole slide images (WSIs) to be exact replicas of the glass slides. All images are scanned at 20x magnification and using the same gain, brightness and exposure times to avoid image to image variations.

The quantitative pathological assessment for the WSIs were analyzed using the HALO® v.3.4.2986 (Indica labs, Albuquerque, New Mexico, USA). For all regions, relevant fields of view were manually traced. For MTG and BA9, layers were segmented using DenseNet194, a deep learning convolutional neural network that is minimally pretrained classifier developed to recognize patterns in the tissue structure provided by Halo. Additional training data was created by manually annotating cortical layers labelled with NeuN in 10 cases. Based on the cellular architecture and the relative position withing the cortical ribbon the following layers were annotated: Layer1 (molecular layer), layer 2 (external granular layer), layer 3 (external pyramidal layer), layer 4 (Internal granular layer) and layers 5-6 (internal pyramidal and multiform layers). Then the trained classifier was applied to the NeuN-labelled sides from all donors. All results of the automatic segmentation were examined by a scientist trained in cortical neuroanatomy and adjusted when necessary. Manual adjustment of the annotations also included removal of staining artifacts and non-parenchymal structures, such as large blood vessels by drawing exclusion areas around them.

Second, using the Serial Section registration tool, all 5 WSIs belonging to the same case (labelled with NeuN, GFAP, α-Syn, Aβ combined with Iba1, and pTau combined with pTDP-43) were registered to each other in order to establish anatomical correspondence between all 5 tissue sections, and the cortical annotations from the NeuN-labelled slide were copied to the other 4 IHC stained slides (noted above). We then applied different algorithms and approaches to obtain stain-specific metrics from all the slides for each cortical layer. Area quantification algorithm (Area Quantification module) was used for determining the area of positive staining for all proteins of interest (NeuN, GFAP, Iba1, Aβ, pTau, α-Syn, and pTDP-43). Multiplex IHC module for used to determine the number of cells displaying positive labelling for NeuN, pTau, α-Syn, and pTDP-43). For the double labelled slides, Multiplex IHC module was used to estimate the area of co-localization of pTau with pTDP-43, and Aβ with Iba1. Microglia Activation module was used to determine the number of cells positive for Iba1, measure the cell process area and length, as well as to classify the cells according to the activation state (activated vs not, based on the process area and thickness). In the slides double-labelled for Aβ and Iba1 the Object Colocalization module was used to determine the number of Aβ-positive objects (amyloid plaques), the average object area, median object diameter, and the number of objects that were double-positive for Aβ and Iba1.

#### Tissue processing for single nucleus isolations

Brain regions were identified on tissue slab photographs taken at the time of autopsy and at the time of dissection using the Allen Human Reference Atlas as a guide for region localization. Caudate nucleus blocks were taken from the A-P level anterior to the appearance of the Globus Pallidus. To dissect regions of interest, tissue slabs were removed from storage at –80C, briefly transferred to a –20C freezer to prevent tissue shattering during dissection, and then handled on a custom cold table maintained at –20C during dissection. Dissections were performed using dry ice-cooled razor blades or scalpels to prevent warming of tissues. Photographs were taken before and after each dissection to document the precise location of each resected tissue block. Dissected tissue samples were then transferred to vacuum seal bags, sealed, and stored at -80C until the time of use. Single nucleus suspensions were generated using a previously described standard procedure (https://www.protocols.io/view/isolation-of-nuclei-from-adult-human-brain-tissue-ewov149p7vr2/v2). Briefly, after tissue homogenization, isolated nuclei were stained with a primary antibody against NeuN (FCMAB317PE, Millipore-Sigma) to label neuronal nuclei. Nuclei samples were analyzed using a BD FACS Aria flow cytometer and nuclei were sorted using a standard gating strategy to exclude multiplets (^107^A defined mixture of neuronal (70% from the NeuN positive gate) and non-neuronal (30% from the NeuN negative) nuclei was sorted for each sample. Nuclei isolated for 10x Genomics v3.1 snRNA-seq were concentrated by centrifugation after FANS and were frozen and stored at –80C until later chip loading. Nuclei isolated for 10x Genomics Multiome and 10x Genomics Single Cell ATAC v1.1 were concentrated by centrifugation after FANS and were immediately processed for chip loading.

#### 10x Genomics sample processing

10x Genomics chip loading and post-processing of the emulsions to sequencing libraries were done with the Chromium Next GEM Single Cell 3’ Gene Expression v3.1, Chromium Next GEM Single Cell ATAC v1.1, and Chromium Next GEM Single Cell Multiome ATAC+Gene Expression kits according to the manufacturer’s guidelines. Nuclei concentration was calculated either manually using a disposable hemocytometer (InCyto, DHC-NO1) or using the NC3000 NucleoCounter.

#### 10x sequencing, demultiplexing, mapping to reference genome

All 10x libraries were sequenced per manufacturer’s specifications on a NovaSeq 6000 using either a NovaSeq-X or S4 flow cell. Reads were demultiplexed to fastq files using BCL Convert (version 4.2.7) for libraries run on NovaSeq-X flow cells and bcl2fastq (version 2-20-0) for libraries run on S4 flow cells. Reads from snRNA-seq libraries were mapped to 10x Genomics’ official human reference (“Human reference (GRCh38) – 2020-A”) and unique molecular identifiers (UMIs) counted per gene using the cellranger (version 6.1.1) pipeline with the “—include—introns” parameter included. Reads from snATAC-seq and snMultiome libraries were mapped to the same reference using cellranger-atac (version 2.0.0) and cellranger-arc (version 2.0.0) pipelines with default parameters, respectively.

#### Xenium methods

Fresh frozen tissue sections were mounted onto Xenium slides (10X Genomics) and stored at −80 °C until use. Slides were equilibrated at 37 °C for 1 min using a pre-heated thermal cycler (Bio-Rad 1851197) equipped with a Xenium Thermocycler Adaptor. Fixation was performed in 1× PBS containing 2.5 mL paraformaldehyde (Electron Microscopy Sciences 15710) for 30 min at room temperature. Slides were permeabilized with 1% SDS (Millipore Sigma 71736) and incubated in pre-chilled 70% methanol for 60 min on wet ice.

Following permeabilization, slides were washed and transferred into Xenium cassettes. Probe hybridization was performed using a mix of Xenium Probe Hybridization Buffer, Probe Dilution Buffer, and either pre-designed or custom gene expression probes (10X Genomics). Probes were preheated at 95 °C for 2 min and cooled on ice prior to mixing. Each slide received 500 µL of hybridization mix and was incubated overnight (16–24 h) at 50 °C. Following hybridization, slides underwent washes using Xenium Post Hybridization Wash Buffer at 37 °C. Ligation was performed by adding 500 µL of freshly prepared ligation mix containing Xenium Ligation Buffer, Enzyme A, and Enzyme B, followed by a 2 h incubation at 37 °C. Amplification was conducted using a master mix of Xenium Amplification Mix and Enzyme, incubated for 2 h at 30 °C. Autofluorescence quenching was achieved by sequential washes in TE buffer, 1× PBS, and ethanol solutions, followed by incubation with Xenium Autofluorescence Solution for 10 min in the dark. Nuclei staining was performed using Xenium Nuclei Staining Buffer, followed by four washes in PBS-T. Slides were stored in PBS-T at 4 °C or immediately processed for imaging.

Prepared slides were loaded into the Xenium Analyzer (10X Genomics 1000481) along with decoding reagent modules, buffer bottles, and consumables. Imaging buffers were freshly prepared, including Xenium Probe Removal Buffer and Sample Wash Buffers A and B. Reagents were loaded into designated positions on the instrument, and system checks were performed prior to initiating the run. Following the run, slides were scanned for DAPI and autofluorescence signals, and imaging regions were designated using the instrument’s touchscreen interface. Imaging runs were initiated and completed over 1–3 days. Upon completion, fluidics cleanup was performed, and slides were stored in PBS-T at 4 °C. Imaging data were exported and reviewed for quality control. All slides processed through Xenium were segmented using the Xenium segmentation kit add-on.

#### Post-Xenium Immunofluorescence Staining

For post-hoc immunofluorescence, Xenium-procesed slides were placed in a staining tray and 500ul of DAPI solution (5ug/mL in PBS) was applied for 15 min at RT in the dark. Slides were then washed 3 x 5 min in 1X TBST in coplin jars with the Corning LSE Platform Rocker set to 120 rpm. Slides were post-fixed with 4% PFA for 30 min, then washed in 1X TBST for 3 x 5 min. Slides were then subjected to antigen retrieval with Diva Decloaker (Biocare Medical, cat# DV2004) using a decloaking chamber Decloaking Chamber™ NxGen (Biocare Medical DC 2012) according to the manufacturer’s instruction, at 90°C for 20 minutes. After cooling for 3+ minutes at RT, slides were washed 3 x 5 min in 1X TBST. Defatting and autofluorescence removal was performed with 10% 2-methyl beta cyclodextrin (2MBCD) dissolved in 1X PBS (pH7.4) for two hours at RT, then washed in 1X TBST for 3x5 min. To block endogenous peroxidase in the tissue, a 1X solution of H_2_O_2_ was prepared by diluting 100X H_2_O_2_ solution (component C2 in Life Technologies, B40915 or B40912 Tyramide SuperBoost Kits) in sufficient volume to cover the selected slides. 500ul of this 1X H_2_O_2_ blocking solution was applied to each slide and incubated at RT in a humidified staining tray chamber for 20 min. Peroxide blocking solution was removed by gently tapping the slides against the staining tray. Slides were washed with 1X TBST for 3 x 5 min at RT, then 500ul blocking solution (10% Goat Serum in 1x TBST) was added to each slide for 1 hr.

Following the goat serum blocking step, 250ul of carrier solution (10% Goat Serum/1xTBST) containing a 1:200 dilution of mouse monoclonal anti-AT8 antibody (Invitrogen, MN1020) was added to each slide and the staining solution was covered with a piece of parafilm, ensuring no bubbles formed underneath. The humidified tray was covered and incubated overnight at 4°C. The parafilm was removed gently to protect tissue, the primary antibody solution was removed by gently tapping against the tray and the slides were washed 3 x 5 min at RT with 1X TBST. The first secondary antibody – goat anti-mouse IgG-HRP conjugate antibody - was prepared and diluted according to the instructions of the Alexa Fluor 488 Tyramide SuperBoost Kit (Life Technologies, B40912), applied to the slides and incubated for 1 hr at RT. Slides were rinsed with 1X TBST. Tyramide signal amplification was performed according to the manufacturers instructions, by preparing a 1x solution of tyramide, H_2_O_2_ and reaction buffer, 250 ul of this solution was applied to each slide and incubated for 7 minutes. This solution was quickly removed and 100 ul Stopping Solution (prepared according to the kit manufacturer’s instructions) was applied to the slide for 30 seconds, followed by 3x5 min TBST washes in a coplin jar covered with tin foil to block light. 50 ml of elution buffer (10% SDS, 25 mM glycine, pH2) pre-heated to 60°C was added to a coplin jar, and the washed slides were added. The jar was covered with tin foil, and the mini-incubator shaker (Fisherbrand, 02-217-753) was set to 60°C to 230 rpm and the slides were incubated for 3.5 hours. Slides were washed 4 x 5 min in TBST at RT. Slides were then blocked with Goat Serum Blocking Solution provided with the SuperBoost Kit for 1 hr at RT. A 1:200 dilution of 6E10 mouse-anti-b-Amyloid,1-16 (Biolegend #803003) was prepared in blocking solution and 250 ul was applied to each slide prior to incubation overnight at 4°C. slides were washed 3 x 5 min in TBST at RT, then the Goat anti-Mouse IgG provided with the AF-594 TSA kit (Life Technologies, B40915) was diluted and 250 ul was added to the slides according to the manufacturer’s instructions. The slides were incubated in a staining tray protected from light for 1 hr at RT. Slides were rinsed with 1X TBST. Tyramide signal amplification was performed according to the manufacturers instructions for the 594 kit, by preparing a 1x solution of tyramide, H_2_O_2_ and reaction buffer, 250 ul of this solution was applied to each slide and incubated for 6 min. This solution was quickly removed and 100 ul Stopping Solution (prepared according to the kit manufacturer’s instructions) was applied to the slide for 30 seconds, followed by 3x5 min TBST washes in a coplin jar covered with tin foil to block light. Slides were dried overnight in the dark in a protected chamber. 200 ul of Prolong Diamond with DAPI (ThermoFisher, cat# P36971 or equivalent) was applied to each slide, the slides were coverslipped in preparation for imaging. Slides were imaged using a VS200 fluorescent slide scanner according to the manufacturer’s instructions.

#### Spatial transcriptomics gene panel selection

An initial set of 300 genes were selected based on the human BG panel selection from Hewitt and Turner et al to be published concurrently^9^. Due to concerns that it was highly expressed across the Ca outside of cell bodeis making cell typing more difficult, *PTP1R1B* was removed from the panel. Addtionally, as *SLC17A7* is primarily a excitatory neuronal marker, we found that it was of limited use in a Ca panel and may confuse cell type mapping algorithms, it was removed. An additional set of 11 genes were added to improve discrimination of MSN subtypes in Ca using NSForest^108^. The remaining 150 genes were a collection of AD related genes collected from the literature. The list of genes in the gene panel are available in supplementary table X.

### QUANTIFICATION AND STATISTICAL ANALYSIS

#### Quantification of Neuropathology

A custom classifier (HALO AI, DenseNet) was developed to assign all Aβ plaques to 3 subtypes: diffuse, fibrillar, or dense core. The training set included 157 examples of diffuse plaques, 254 fibrillar, and 87 dense core plaques. Another custom classifier (HALO AI, DenseNet) was trained to recognize neurofibrillary tangles in sections labeled with pTau antibody. The classifier was trained on samples of cerebral cortex (7179 examples) and the hippocampus (5770 examples). Development, optimization, and testing of all analysis algorithms was done by a scientist trained in neuropathology. The final quantitative neuropathology dataset includes raw measurements (absolute values) and metrics normalized to the unit area. All quantifications are available in Supplemental Table 2.

#### Creating a combined reference atlas of SEA-AD and HMBA for snRNA-seq and Spatial Transcriptomics

We integrated the snRNA-seq SEA-AD MTG reference taxonomy (reference) with the Human Ca (head and body) portion of the HMBA BG cross species reference taxonomy (query) utilizing iterative scANVI as described in Gabitto and Travaglini et al 2024^5^. For the HMBA taxonomy, we only utilized cells sampled from human and in the Ca head or body. The latent space corresponding to neuron vs. non-neuron classification was then clustered using de novo clustering with the *scanpy.tl.leiden* function^109^. To measure the overlap between the two atlases per cluster, we utilize the *scipy.stats.entropy* to calculate the mixing per cell based on its nearest neighbors, and then taking the mean value per cluster^110^. We set an overlap threshold at 0.075, where cells from the HMBA reference atlas in clusters above said threshold were labeled using their new SEA-AD labels, while clusters below said threshold kept their HMBA label. This relabeled combined atlas was used as reference to query the labels for SEA-AD Ca cells when using iterative scANVI.

For quality control (QC) we first filtered cells utilizing initial heuristics where we keep all cells with above 2000 and 100000 unique molecular identifiers (UMIs). Additionally we kept all cells that had between 1000 and 13000 genes detected. All cells with a doublet score of less than 0.3 were kept. All cells with a fraction of mitochondrial UMIs under 0.03 were kept. For all neuronal cells, an additional QC heuristic was applied where we only kept neurons with at least 2000 genes. QC was also applied on a per cell basis based on their cell type specific latent space. Areas of the nearest neighbor graph on the latent space with few reference nuclei could represent droplets with ambient RNA, multiplet nuclei, dying cells, or transcriptional states missing from the reference, unique to a donor, or found only in aging or disease. To assess these possibilities, we fractured the graph into tens to hundreds of clusters using high resolution Leiden clustering implemented in *the scanpy.tl.leiden* function. Clusters were then flagged and removed if they had poor group doublet scores^107^, fraction of mitochondrial reads, number of genes detected, with cutoffs adjusted for each cell type at each level of the hierarchy based on their distributions (to account for dramatically different RNA content found across cell types).

For QC in spatial transcriptomics we note that the majority of cells were segmented primarily using a combination of the DAPI and the 18S RNA interior stain from the 10x segmentation kit. The boundary stain (ATP1A1/E-cadherin/CD45) proved too blown out for clean discrimination between cell bodies likely due to the dense and entangled nature of the neuropil. The interior proteins stain (alphaSMA/Vimentin) mostly was used to segment cells near where sections were cut and we expect to find poor quality cells. For these reasons, only cells segmented with 18S RNA interior stain were kept. We considered cells with less than 15 genes and 30 transcripts (from a lack of hybridization, due to the cell being out of plane, etc.) to be unmappable and thus were filtered from the dataset. All of these cells were then integrated utilizing iterative scANVI with several modifications. One, the gene set was fixed to the original 461 genes of the gene panel at each level of the taxonomic hierarchy. Two, the latent space dimensionality was set to 30 rather than 10. Third, the classifier for scANVI was set to linear as a way to improve stability in the integration. The authors note that measurement error and noise in spatial transcriptomics, especially that from segmentation error remains an open problem^111^. As such the integration remains imperfect and thus the heuristic of out of distribution cells on the latent space potentially representing poor quality cells no longer holds as strongly. As a result, we elected to not perform per cell type filtering using Leiden clustering on the latent space as described before for snRNA-seq.

#### Modeling Disease Progression in the Caudate Nucleus

To infer a continuous measure of Alzheimer’s disease progression in the Ca, we developed a hierarchical Bayesian model that jointly estimated latent pseudotimes for pTau and amyloid Aβ pathologies. Four quantitative neuropathology measurements were used per donor: (1) number of AT8-positive cells per area, (2) percent AT8-positive area, (3) number of 6E10-positive objects per area, and (4) percent 6E10-positive area (Supplemental Table 2). Pathology features were scaled and ordered by the first principal component across donors to initialize pseudotime inference.

Each donor, *d*, was assigned a central latent pseudotime 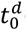 describing CPS, drawn from a Beta distribution with LogNormal hyperpriors on its shape parameters:

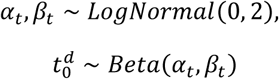

AT8 and 6E10 -specific CPS, 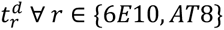, were modeled as Gaussian perturbations around the central pseudotime with variance, 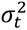:

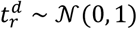

Each observed feature 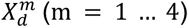 was modeled using a Negative Binomial likelihood with a logit link:

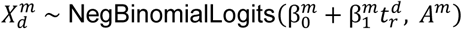

where 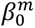 and 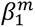 capture the baseline and temporal progression dynamics of feature *m*, and *A*^*m*^∼*HalfNormal*(50) represents feature-specific dispersion. A fixed assignment matrix linked each feature to its corresponding pathology class (*AT*8 *features* → *AT*8, 6*E*10 *features* → 6*E*10), reducing the number of free parameters and improving identifiability.

The model was implemented in NumPyro using the No-U-Turn Sampler (NUTS). Sampling was performed with 10,000 warmup and 1,000 posterior draws, repeated across five random seeds. Inference outputs were summarized using ArviZ^112,113^. Convergence was verified with 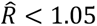 and effective sample size ≫ 50 for the overall disease progression parameter 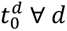. Posterior samples of 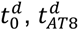, and 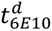 were rescaled to [0,1] to obtain the final continuous pseudoprogression scores. Donor CPS values are available in Supplemental Table 7.

#### Differential abundance analysis

For differential analyses we filtered all cell types with fewer than ten cells across all library preparations. To model changes in the composition of cell types as a function of CPS, we used the Bayesian method scCODA as implemented in pertpy^50,114^. We tested compositional changes in neuronal and non-neuronal nuclei separately because they were sorted to have a defined ratio (70% neuronal nuclei, 30% nonneuronal nuclei in each donor). To do this, we created separate AnnData objects of neuronal and nonneuronal nuclei with cell-type annotations, sequencing library IDs and relevant donor-level covariate information. As we did not know which supertypes would be affected by AD, we ran models with each supertype set as the unchanged reference population, as recommended by the authors of scCODA. We set up an ensemble of models to test whether supertypes were credibly affected across CPS (interval (0,1)) using the pertpy’s *Sccoda* class function *run_nuts* with formula set to sex + age at death + 10X chemistry + *APOE4* status + CPS_x_ ∀ *x* ∈ {*Global* (0), 6*E*10, *AT*8}. Here *APOE*4 *Status* was set to 1 if the donor for both homozygous and heterozygous *APOE*4 donors and 0 otherwise. Using each cell type representing at least 4% of the population of cells as the reference celltype, we obtained 1000 posterior estimates for each parameter after 1000 warmup. The sampling occasionally stayed at fixed points, so we reran models with fewer than 60% accepted epochs or more than 95% epochs. We defined credibly affected supertypes as those that had a mean inclusion probability across models greater than 0.85. All differential abundances are available in Supplemental Table 3.

#### Connecting microglial factors to microglial markers

Microglial marker differential expression was taken from Gabitto and Travaglini et al where they performed differential expression on one vs. all cell types comparison in their dataset^5^. Microglial factors were taken from Marshe et al, and were converted into gene sets by taking the genes with the top 100 factor loadings^57^. We ranked genes from the marker gene differential expression by calculating *β*_*comparison*_ ∗ − log(*p*_*adj*_) and then used the *prerank* function from GSEApy^63^. All factors with an adjusted p < 0.05 was determined to be significant.

#### Differential Expression Analysis

To model gene expression changes along CPS while adjusting for other covariates and pseudo-replication within donors we used a general linear mixed effects model implemented in the NEBULA R package which we accessed in python with the rpy2 package^115^. For snRNA-seq differential expression the formula was set to *y* ∼ *Age* + *Sex* + *APOE*4 *Status* + 10*X Chemistry* + *CPS*_*x*_ ∀ *x* ∈ {*Global* (0), 6*E*10, *AT*8}. Here *APOE*4 *Status* was set to 1 if the donor for both homozygous and heterozygous *APOE*4 donors and 0 otherwise. The sample term was set to the donor ID. The intercept term was set to the number of transcripts counted in a cell. All genes with mean counts across all cells less than 0.005 were removed from the differential expression analysis. All genes that did not converge according to NEBULA or had an overdispersion of above 1 were removed. For each gene, each *β*_*x*_with a fitting standard error of less than 0.05 and greater than 0.5 were also removed due to potential general linear mixed effects model convergence issues not detected by NEBULA. All p values were adjusted using the Benjamani-Hochberg method. All genes with an adjusted *p*_*CPSx*_ < 0.1 and a |*β*_*CPSx*_ | > 0.5 were significant. All differential expression results are available in Supplemental Tables 4 – 6.

#### Normalizing transcript counts for visualization

For both the spatial and single nucleus transcriptomics the shifted log method was utilized to normalize counts^116^. The basic function on the counts in modalities utilized was:

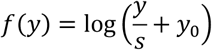

Where *y* is the UMIs, s being a so-called size factor and *y*_0_ describing a pseudo-count which we set to 1. The size factor, *s*, which is applied to account for variations across cells is defined in snRNA-seq as:

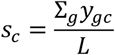

Where *c* is indexing cells, and *g* is indexing genes. *L* is the target sum which we set equal to 10000. In spatial transcriptomics instead of using the natural log, we use log base-2 function and *s* is just the volume of the segmentation polygon of the cell.

#### Processing post-Xenium immunofluorescence staining

Alignment between the spatial transcriptomics and immunofluorescence images was performed on the DAPI signal using Xenium Explorer software^117^. After alignment, each section’s xenium files and aligned post-hoc immunofluorescence image was summarized into *spatialdata* zarr files^118^. Aligned immunofluoresence images were composed of a DAPI channel, an AT8 channel, and a 6E10 channel. To classify pixels that were pTau positive in the AT8 channel of the image, we applied a difference of gaussian filter utilizing the python *skimage.filters* module with low_sigma being 5 mm, and high sigma being at big sigma being at 20 mm^119^. All signal 3 standard deviations above the mean was kept. To classify pixels in that were A*β* positive in the 6E10 channel of the image, we took pixels with signal 3 standard deviations from the mean and then applied erosion with a 3x1 footprint on the binary mask utilizing *skimage.morphology.binary_erosion*. Holes in the label mask were removed utilizing the *skimage.morpholgy.remove_small_holes* function. Both the pTau and A*β* images were converted to labels using *skimage.morphology.labels* or its *dask.ndmeasure* counterpart. Label objects smaller than 64 pixels in area (0.2125 mm per pixel) were filtered.

To remove autofluorescence contamination from lipofusin in the image channel corresponding to AT8, non-negative matrix factorization (NMF) with three factors (corresponding to the AT8 signal, 6E10 signal, and the autofluorescence signal) was done on the AT8 and the 6E10 channels of each post-hoc immunofluorescence image. NMF was performed on the image using the *sklearn.decomposition.NMF*^120^. We note that NMF component weights for the lipofusin channel are ∼1:20, 6E10 to AT8. Thus, unmixing was done by calculating the cosine similarity of each pixel vector to each NMF component and then mask pTau label pixels that didn’t have a higher similarity to the AT8 component than the autofluorescence component, or had a higher similarity to the vector [1,1] than the autofluorescence component.

Calculating pTau overlap with cell bodies nuclei was done utilizing the masked pTau labels. Both cell body labels and nuclei labels were taken from the xenium segmentation output. Here areas classified to contain pTau in the image were set to 1 and the rest were set to 0. In both cases, *spatialdata.aggregate* with the *agg_funct=’mean’* was used to measure spatial overlap between pTau and cells.

#### Binning regions for white matter and pTau analysis

We binned Xenium sections into 2500 µm bins and calculated oligodendrocyte area and MSN neuronal area in each bin. To estimate the enrichment in regions, bins were separated into white matter dense (oligodendrocyte coverage was greater than or equal to 10% of the bin area) and white matter sparse regions (oligodendrocyte coverage was less than 10% bin area). pTau coverage was normalized by the area of the bin. Significance was determined by the Wilcoxon rank-sum test. The same was done for microglial coverage.

#### Neighborhood Enrichment Analysis

We take segmentation polygons for cells and pathology, and then calculate all neighbors within 40 *μ*m of a cell using the sort-tile-recursive algorithm implemented in *shapely.STRtree*^121^. Within those set of cells, we calculate the minimum distance between polygons. We measure the number of times cells a particular type were observed within 5 *μ*m bins from the polygon of interest (join counts). A null distribution was produced by randomly translating cell and pathology polygons 1000 times within 100 *μ*m of their original location. A Z-score (neighborhood enrichment score) and p-value was calculated using the observed join counts and the join counts from the null distribution. The p-value was adjusted using the Benjamini-Hochberg method, and all enrichment scores with an adjusted p-value less than 0.05 were set to 0. We considered all scores > 3 (representing 3 standard deviations from the null mean) to be significant

## ADDITIONAL RESOURCES

Study data are freely available at the SEA-AD’s website: https://brain-map.org/consortia/sea-ad. Access to specific ACT Study data may require execution of appropriate human subjects review and data sharing agreements by the process described on the Adult Changes in Thought (ACT) website: actagingresearch.org.

## REFERENCES

1. Yao, Z., Van Velthoven, C.T.J., Kunst, M., Zhang, M., McMillen, D., Lee, C., Jung, W., Goldy, J., Abdelhak, A., Baker, P., et al. (2023). A high-resolution transcriptomic and spatial atlas of cell types in the whole mouse brain (Neuroscience) 10.1101/2023.03.06.531121.

2. Johansen, N., Fu, Y., Schmitz, M., and, et al. (2025). Cross-Species Consensus Atlas of the Primate Basal Ganglia. *in preparation*.

3. BRAIN Initiative Cell Census Network (BICCN), BRAIN Initiative Cell Census Network (BICCN) Corresponding authors, Callaway, E.M., Dong, H.-W., Ecker, J.R., Hawrylycz, M.J., Huang, Z.J., Lein, E.S., Ngai, J., Osten, P., et al. (2021). A multimodal cell census and atlas of the mammalian primary motor cortex. Nature 598, 86–102. 10.1038/s41586-021-03950-0.

4. Siletti, K., Hodge, R., Mossi Albiach, A., Lee, K.W., Ding, S.-L., Hu, L., Lönnerberg, P., Bakken, T., Casper, T., Clark, M., et al. (2023). Transcriptomic diversity of cell types across the adult human brain. Science 382, eadd7046. 10.1126/science.add7046.

5. Gabitto, M.I., Travaglini, K.J., Rachleff, V.M., Kaplan, E.S., Long, B., Ariza, J., Ding, Y., Mahoney, J.T., Dee, N., Goldy, J., et al. (2024). Integrated multimodal cell atlas of Alzheimer’s disease. Nat Neurosci 27, 2366–2383. 10.1038/s41593-024-01774-5.

6. Hawrylycz, M., Kaplan, E.S., Travaglini, K.J., Gabitto, M.I., Miller, J.A., Ng, L., Close, J.L., Hodge, R.D., Long, B., Mollenkopf, T., et al. (2024). SEA-AD is a multimodal cellular atlas and resource for Alzheimer’s disease. Nat Aging 4, 1331–1334. 10.1038/s43587-024-00719-8.

7. BICAN Consortium (2025). A Community Standard Multispecies Cell Atlas of the Basal Ganglia: The BRAIN Initiative Cell Atlas Network. *in preparation*.

8. Lui, X.-P. and, et al. (2025). Morphoelectric Diversity and Specialization of Neuronal Cell Types in the Primate Striatum. *in preperation*.

9. Hewitt, M.N., Turner, M.A., Johansen, N., McMillen, D.A., Dan, S., DeBerardine, M., Ruiz, A., Huang, M., Quon, J., Fu, Y., et al. (2025). A cross-species spatial transcriptomic atlas of the human and non-human primate basal ganglia. Preprint, 10.1101/2025.11.22.688128 https://doi.org/10.1101/2025.11.22.688128.

10. Graybiel, A.M. (2000). The basal ganglia. Current Biology 10, R509–R511. 10.1016/S0960-9822(00)00593-5.

11. Morgenstern, N.A., Isidro, A.F., Israely, I., and Costa, R.M. (2022). Pyramidal tract neurons drive amplification of excitatory inputs to striatum through cholinergic interneurons. Sci. Adv. 8, eabh4315. 10.1126/sciadv.abh4315.

12. Myers, R.H., Vonsattel, J.P., Paskevich, P.A., Kiely, D.K., Stevens, T.J., Cupples, L.A., Richardson, E.P., and Bird, E.D. (1991). Decreased neuronal and increased oligodendroglial densities in Huntington’s disease caudate nucleus. J Neuropathol Exp Neurol 50, 729–742. 10.1097/00005072-199111000-00005.

13. Mandelli, M.L., Savoiardo, M., Minati, L., Mariotti, C., Aquino, D., Erbetta, A., Genitrini, S., Di Donato, S., Bruzzone, M.G., and Grisoli, M. (2010). Decreased Diffusivity in the Caudate Nucleus of Presymptomatic Huntington Disease Gene Carriers: Which Explanation? AJNR Am J Neuroradiol 31, 706–710. 10.3174/ajnr.a1891.

14. Lach, B., Grimes, D., Benoit, B., and Minkiewicz-Janda, A. (1992). Caudate nucleus pathology in Parkinson’s disease: ultrastructural and biochemical findings in biopsy material. Acta Neuropathol 83, 352–360. 10.1007/BF00713525.

15. 15. De Simoni, S., Jenkins, P.O., Bourke, N.J., Fleminger, J.J., Hellyer, P.J., Jolly, A.E., Patel, M.C., Cole, J.H., Leech, R., and Sharp, D.J. (2018). Altered caudate connectivity is associated with executive dysfunction after traumatic brain injury. Brain 141, 148–164. 10.1093/brain/awx309.

16. Xu, H., Zhang, X., and Bai, G. (2022). Abnormal Dorsal Caudate Activation Mediated Impaired Cognitive Flexibility in Mild Traumatic Brain Injury. J Clin Med 11, 2484. 10.3390/jcm11092484.

17. Wylie, G.R., Dobryakova, E., DeLuca, J., Chiaravalloti, N., Essad, K., and Genova, H. (2017). Cognitive fatigue in individuals with traumatic brain injury is associated with caudate activation. Sci Rep 7, 8973. 10.1038/s41598-017-08846-6.

18. Marié, R.M., Barré, L., Dupuy, B., Viader, F., Defer, G., and Baron, J.C. (1999). Relationships between striatal dopamine denervation and frontal executive tests in Parkinson’s disease. Neuroscience Letters 260, 77–80. 10.1016/s0304-3940(98)00928-8.

19. Rinne, J.O., Portin, R., Ruottinen, H., Nurmi, E., Bergman, J., Haaparanta, M., and Solin, O. (2000). Cognitive Impairment and the Brain Dopaminergic System in Parkinson Disease: [^18^F]Fluorodopa Positron Emission Tomographic Study. Arch Neurol 57, 470. 10.1001/archneur.57.4.470.

20. 20. Müller, U., Wächter, T., Barthel, H., Reuter, M., and Von Cramon, D.Y. (2000). Striatal [ 123 I]β-CIT SPECT and prefrontal cognitive functions in Parkinson’s disease. Journal of Neural Transmission 107, 303–319. 10.1007/s007020050025.

21. 21. Nobili, F., Campus, C., Arnaldi, D., De Carli, F., Cabassi, G., Brugnolo, A., Dessi, B., Morbelli, S., Sambuceti, G., Abbruzzese, G., et al. (2010). Cognitive-nigrostriatal relationships in de novo, drug-naïve Parkinson’s disease patients: a [I-123]FP-CIT SPECT study. Mov Disord 25, 35–43. 10.1002/mds.22899.

22. Luquez, T., Algoo, J., Chiu, R., Mares, J.A., Yadav, A., Lam, M., Gaur, P., Lai, X., Lee, D.I., Paryani, F., et al. (2025). Cell type-specific associations with Alzheimer’s Disease conserved across racial and ethnic groups. Preprint at Neuroscience, 10.1101/2025.04.14.648597 https://doi.org/10.1101/2025.04.14.648597.

23. Thal, D.R., Rub, U., Orantes, M., and Braak, H. (2002). Phases of A beta-deposition in the human brain and its relevance for the development of AD. Neurology 58, 1791–1800.

24. Matsushima, A., Pineda, S.S., Crittenden, J.R., Lee, H., Galani, K., Mantero, J., Tombaugh, G., Kellis, M., Heiman, M., and Graybiel, A.M. (2023). Transcriptional vulnerabilities of striatal neurons in human and rodent models of Huntington’s disease. Nat Commun 14, 282. 10.1038/s41467-022-35752-x.

25. Villalba, R.M., and Smith, Y. (2013). Differential striatal spine pathology in Parkinson’s disease and cocaine addiction: a key role of dopamine? Neuroscience 251, 2–20. 10.1016/j.neuroscience.2013.07.011.

26. Zaja-Milatovic, S., Milatovic, D., Schantz, A.M., Zhang, J., Montine, K.S., Samii, A., Deutch, A.Y., and Montine, T.J. (2005). Dendritic degeneration in neostriatal medium spiny neurons in Parkinson disease. Neurology 64, 545–547. 10.1212/01.WNL.0000150591.33787.A4.

27. Stephens, B., Mueller, A.J., Shering, A.F., Hood, S.H., Taggart, P., Arbuthnott, G.W., Bell, J.E., Kilford, L., Kingsbury, A.E., Daniel, S.E., et al. (2005). Evidence of a breakdown of corticostriatal connections in Parkinson’s disease. Neuroscience 132, 741–754. 10.1016/j.neuroscience.2005.01.007.

28. Liang, L., DeLong, M.R., and Papa, S.M. (2008). Inversion of Dopamine Responses in Striatal Medium Spiny Neurons and Involuntary Movements. J. Neurosci. 28, 7537–7547. 10.1523/JNEUROSCI.1176-08.2008.

29. Taverna, S., Ilijic, E., and Surmeier, D.J. (2008). Recurrent Collateral Connections of Striatal Medium Spiny Neurons Are Disrupted in Models of Parkinson’s Disease. J. Neurosci. 28, 5504–5512. 10.1523/JNEUROSCI.5493-07.2008.

30. Jellinger, K.A., and Attems, J. (2006). Does striatal pathology distinguish Parkinson disease with dementia and dementia with Lewy bodies? Acta Neuropathol 112, 253–260. 10.1007/s00401-006-0088-2.

31. Espay, A.J., Vizcarra, J.A., Marsili, L., Lang, A.E., Simon, D.K., Merola, A., Josephs, K.A., Fasano, A., Morgante, F., Savica, R., et al. (2019). Revisiting protein aggregation as pathogenic in sporadic Parkinson and Alzheimer diseases. Neurology 92, 329–337. 10.1212/WNL.0000000000006926.

32. Davis, M.Y., Keene, C.D., Jayadev, S., and Bird, T. (2014). The Co-Occurrence of Alzheimer’s Disease and Huntington’s Disease: A Neuropathological Study of 15 Elderly Huntington’s Disease Subjects. Journal of Huntington’s Disease 3, 209–217. 10.3233/JHD-140111.

33. St-Amour, I., Turgeon, A., Goupil, C., Planel, E., and Hébert, S.S. (2018). Co-occurrence of mixed proteinopathies in late-stage Huntington’s disease. Acta Neuropathol 135, 249–265. 10.1007/s00401-017-1786-7.

34. 2017 Alzheimer’s disease facts and figures (2017). Alzheimer’s & Dementia 13, 325–373. 10.1016/j.jalz.2017.02.001.

35. Keren-Shaul, H., Spinrad, A., Weiner, A., Matcovitch-Natan, O., Dvir-Szternfeld, R., Ulland, T.K., David, E., Baruch, K., Lara-Astaiso, D., Toth, B., et al. (2017). A Unique Microglia Type Associated with Restricting Development of Alzheimer’s Disease. Cell 169, 1276–1290.e17. 10.1016/j.cell.2017.05.018.

36. Deczkowska, A., Keren-Shaul, H., Weiner, A., Colonna, M., Schwartz, M., and Amit, I. (2018). Disease-Associated Microglia: A Universal Immune Sensor of Neurodegeneration. Cell 173, 1073–1081. 10.1016/J.CELL.2018.05.003.

37. Prater, K.E., Green, K.J., Mamde, S., Sun, W., Cochoit, A., Smith, C.L., Chiou, K.L., Heath, L., Rose, S.E., Wiley, J., et al. (2023). Human microglia show unique transcriptional changes in Alzheimer’s disease. Nat Aging 3, 894–907. 10.1038/s43587-023-00424-y.

38. Probabilistic harmonization and annotation of single-cell transcriptomics data with deep generative models | Molecular Systems Biology https://www.embopress.org/doi/full/10.15252/msb.20209620.

39. McKeith, I.G., Galasko, D., Kosaka, K., Perry, E.K., Dickson, D.W., Hansen, L.A., Salmon, D.P., Lowe, J., Mirra, S.S., Byrne, E.J., et al. (1996). Consensus guidelines for the clinical and pathologic diagnosis of dementia with Lewy bodies (DLB): report of the consortium on DLB international workshop. Neurology 47, 1113–1124.

40. Gearing, M., Levey, A.I., and Mirra, S.S. (1997). Diffuse Plaques in the Striatum in Alzheimer Disease (AD): Relationship to the Striatal Mosaic and Selected Neuropeptide Markers. Journal of Neuropathology and Experimental Neurology 56, 1363–1370. 10.1097/00005072-199712000-00011.

41. Suenaga, T., Hirano, A., Llena, J.F., Yen, S.-H., and Dickson, D.W. (1990). Modified Bielschowsky stain and immunohistochemical studies on striatal plaques in Alzheimer’s disease. Acta Neuropathol 80, 280–286. 10.1007/BF00294646.

42. Masliah, E., Terry, R.D., Mallory, M., Alford, M., and Hansen, L.A. (1990). Diffuse plaques do not accentuate synapse loss in Alzheimer’s disease. Am J Pathol 137, 1293–1297.

43. Liu, F., Sun, J., Wang, X., Jin, S., Sun, F., Wang, T., Yuan, B., Qiu, W., and Ma, C. (2022). Focal-type, but not Diffuse-type, Amyloid Beta Plaques are Correlated with Alzheimer’s Neuropathology, Cognitive Dysfunction, and Neuroinflammation in the Human Hippocampus. Neurosci. Bull. 38, 1125–1138. 10.1007/s12264-022-00927-5.

44. Braak, H., Alafuzoff, I., Arzberger, T., Kretzschmar, H., and Del Tredici, K. (2006). Staging of Alzheimer disease-associated neurofibrillary pathology using paraffin sections and immunocytochemistry. Acta Neuropathologica 112, 389–404. 10.1007/s00401-006-0127-z.

45. De Jong, L.W., Ferrarini, L., Van Der Grond, J., Milles, J.R., Reiber, J.H.C., Westendorp, R.G.J., Bollen, E.L.E.M., Middelkoop, H.A.M., and Van Buchem, M.A. (2011). Shape Abnormalities of the Striatum in Alzheimer’s Disease. Journal of Alzheimer’s Disease 23, 49–59. 10.3233/JAD-2010-101026.

46. Pievani, M., Bocchetta, M., Boccardi, M., Cavedo, E., Bonetti, M., Thompson, P.M., and Frisoni, G.B. (2013). Striatal morphology in early-onset and late-onset Alzheimer’s disease: a preliminary study. Neurobiol Aging 34, 1728–1739. 10.1016/j.neurobiolaging.2013.01.016.

47. Jiji, S., Smitha, K.A., Gupta, A.K., Pillai, V.P.M., and Jayasree, R.S. (2013). Segmentation and volumetric analysis of the caudate nucleus in Alzheimer’s disease. Eur J Radiol 82, 1525–1530. 10.1016/j.ejrad.2013.03.012.

48. Barber, R., McKeith, I., Ballard, C., and O’Brien, J. (2002). Volumetric MRI study of the caudate nucleus in patients with dementia with Lewy bodies, Alzheimer’s disease, and vascular dementia. J Neurol Neurosurg Psychiatry 72, 406–407. 10.1136/jnnp.72.3.406.

49. Agrawal, A., Rachleff, V.M., Travaglini, K.J., Mukherjee, S., Crane, P.K., Hawrylycz, M., Keene, C.D., Lein, E., Mena, G.E., and Gabitto, M.I. (2024). B-BIND: Biophysical Bayesian Inference for Neurodegenerative Dynamics. bioRxiv, 2024.06.10.597236. 10.1101/2024.06.10.597236.

50. Büttner, M., Ostner, J., Müller, C.L., Theis, F.J., and Schubert, B. (2021). scCODA is a Bayesian model for compositional single-cell data analysis. Nature Communications 12, 6876. 10.1038/s41467-021-27150-6.

51. Mathys, H., Peng, Z., Boix, C.A., Victor, M.B., Leary, N., Babu, S., Abdelhady, G., Jiang, X., Ng, A.P., Ghafari, K., et al. (2023). Single-cell atlas reveals correlates of high cognitive function, dementia, and resilience to Alzheimer’s disease pathology. Cell 186, 4365–4385.e27. 10.1016/j.cell.2023.08.039.

52. Green, G.S., Fujita, M., Yang, H.-S., Taga, M., McCabe, C., Cain, A., White, C.C., Schmidtner, A.K., Zeng, L., Wang, Y., et al. (2023). Cellular dynamics across aged human brains uncover a multicellular cascade leading to Alzheimer’s disease (Neuroscience) 10.1101/2023.03.07.531493.

53. Chen, W.-T., Lu, A., Craessaerts, K., Pavie, B., Sala Frigerio, C., Corthout, N., Qian, X., Laláková, J., Kühnemund, M., Voytyuk, I., et al. (2020). Spatial Transcriptomics and In Situ Sequencing to Study Alzheimer’s Disease. Cell 182, 976–991.e19. 10.1016/j.cell.2020.06.038.

54. Moloney, C.M., Lowe, V.J., and Murray, M.E. (2021). Visualization of neurofibrillary tangle maturity in Alzheimer’s disease: A clinicopathologic perspective for biomarker research. Alzheimer’s & Dementia 17, 1554–1574. 10.1002/alz.12321.

55. Zeng, H., Huang, J., Zhou, H., Meilandt, W.J., Dejanovic, B., Zhou, Y., Bohlen, C.J., Lee, S.-H., Ren, J., Liu, A., et al. (2023). Integrative in situ mapping of single-cell transcriptional states and tissue histopathology in a mouse model of Alzheimer’s disease. Nat Neurosci 26, 430–446. 10.1038/s41593-022-01251-x.

56. Sepulcre, J., Schultz, A.P., Sabuncu, M., Gomez-Isla, T., Chhatwal, J., Becker, A., Sperling, R., and Johnson, K.A. (2016). In Vivo Tau, Amyloid, and Gray Matter Profiles in the Aging Brain. Journal of Neuroscience 36, 7364–7374. 10.1523/JNEUROSCI.0639-16.2016.

57. Marshe, V.S., Tuddenham, J.F., Chen, K., Chiu, R., Haage, V.C., Ma, Y., Lee, A.J., Shneider, N.A., Agin-Liebes, J.P., Alcalay, R.N., et al. (2025). A factor-based analysis of individual human microglia uncovers regulators of an Alzheimer-related transcriptional signature. bioRxiv, 2025.03.27.641500. 10.1101/2025.03.27.641500.

58. Mathys, H., Davila-Velderrain, J., Peng, Z., Gao, F., Mohammadi, S., Young, J.Z., Menon, M., He, L., Abdurrob, F., Jiang, X., et al. (2019). Single-cell transcriptomic analysis of Alzheimer’s disease. Nature 570, 332–337. 10.1038/s41586-019-1195-2.

59. Krasemann, S., Madore, C., Cialic, R., Baufeld, C., Calcagno, N., El Fatimy, R., Beckers, L., O’Loughlin, E., Xu, Y., Fanek, Z., et al. (2017). The TREM2-APOE Pathway Drives the Transcriptional Phenotype of Dysfunctional Microglia in Neurodegenerative Diseases. Immunity 47, 566–581.e9. 10.1016/j.immuni.2017.08.008.

60. Ferreira, R., Santos, T., Viegas, M., Cortes, L., Bernardino, L., Vieira, O.V., and Malva, J.O. (2011). Neuropeptide Y inhibits interleukin-1β-induced phagocytosis by microglial cells. J Neuroinflammation 8, 169. 10.1186/1742-2094-8-169.

61. Chen, W.-C., Liu, Y.-B., Liu, W.-F., Zhou, Y.-Y., He, H.-F., and Lin, S. (2020). Neuropeptide Y Is an Immunomodulatory Factor: Direct and Indirect. Front Immunol 11, 580378. 10.3389/fimmu.2020.580378.

62. Yin, Z., Herron, S., Silveira, S., Kleemann, K., Gauthier, C., Mallah, D., Cheng, Y., Margeta, M.A., Pitts, K.M., Barry, J.-L., et al. (2023). Identification of a protective microglial state mediated by miR-155 and interferon-γ signaling in a mouse model of Alzheimer’s disease. Nat Neurosci 26, 1196–1207. 10.1038/s41593-023-01355-y.

63. Fang, Z., Liu, X., and Peltz, G. (2023). GSEApy: a comprehensive package for performing gene set enrichment analysis in Python. Bioinformatics 39, btac757. 10.1093/bioinformatics/btac757.

64. Bengtsson Gonzales, C., Hunt, S., Munoz-Manchado, A.B., McBain, C.J., and Hjerling-Leffler, J. (2020). Intrinsic electrophysiological properties predict variability in morphology and connectivity among striatal Parvalbumin-expressing Pthlh-cells. Sci Rep 10, 15680. 10.1038/s41598-020-72588-1.

65. Muñoz-Manchado, A.B., Bengtsson Gonzales, C., Zeisel, A., Munguba, H., Bekkouche, B., Skene, N.G., Lönnerberg, P., Ryge, J., Harris, K.D., Linnarsson, S., et al. (2018). Diversity of Interneurons in the Dorsal Striatum Revealed by Single-Cell RNA Sequencing and PatchSeq. Cell Reports 24, 2179–2190.e7. 10.1016/j.celrep.2018.07.053.

66. Klug, J.R., Engelhardt, M.D., Cadman, C.N., Li, H., Smith, J.B., Ayala, S., Williams, E.W., Hoffman, H., and Jin, X. (2018). Differential inputs to striatal cholinergic and parvalbumin interneurons imply functional distinctions. eLife 7, e35657. 10.7554/eLife.35657.

67. Bennett, B.D., and Bolam, J.P. (1994). Synaptic input and output of parvalbumin-immunoreactive neurons in the neostriatum of the rat. Neuroscience 62, 707–719. 10.1016/0306-4522(94)90471-5.

68. Tepper, J.M., Tecuapetla, F., Koós, T., and Ibáñez-Sandoval, O. (2010). Heterogeneity and Diversity of Striatal GABAergic Interneurons. Front Neuroanat 4, 150. 10.3389/fnana.2010.00150.

69. Wippold, F.J., Cairns, N., Vo, K., Holtzman, D.M., and Morris, J.C. (2008). Neuropathology for the Neuroradiologist: Plaques and Tangles. AJNR Am J Neuroradiol 29, 18–22. 10.3174/ajnr.A0781.

70. Sheng, J.G., Zhou, X.Q., Mrak, R.E., and Griffin, W.S.T. (1998). Progressive Neuronal Injury Associated with Amyloid Plaque Formation in Alzheimer Disease: Journal of Neuropathology and Experimental Neurology 57, 714–717. 10.1097/00005072-199807000-00008.

71. Tsering, W., Hery, G.P., Phillips, J.L., Lolo, K., Bathe, T., Villareal, J.A., Ruan, I.Y., and Prokop, S. (2023). Transformation of non-neuritic into neuritic plaques during AD progression drives cortical spread of tau pathology via regenerative failure. acta neuropathol commun 11, 190. 10.1186/s40478-023-01688-6.

72. on behalf of the MRC Cognitive Function and Ageing Neuropathology Study Group, Wharton, S.B., Minett, T., Drew, D., Forster, G., Matthews, F., Brayne, C., and Ince, P.G. (2016). Epidemiological pathology of Tau in the ageing brain: application of staging for neuropil threads (BrainNet Europe protocol) to the MRC cognitive function and ageing brain study. acta neuropathol commun 4, 11. 10.1186/s40478-016-0275-x.

73. Mitchell, T.W., Mufson, E.J., Schneider, J.A., Cochran, E.J., Nissanov, J., Han, L.-Y., Bienias, J.L., Lee, V.M.-Y., Trojanowski, J.Q., Bennett, D.A., et al. (2002). Parahippocampal tau pathology in healthy aging, mild cognitive impairment, and early Alzheimer’s disease. Ann Neurol 51, 182–189. 10.1002/ana.10086.

74. Mitchell, T.W., Nissanov, J., Han, L.Y., Mufson, E.J., Schneider, J.A., Cochran, E.J., Bennett, D.A., Lee, V.M., Trojanowski, J.Q., and Arnold, S.E. (2000). Novel method to quantify neuropil threads in brains from elders with or without cognitive impairment. J Histochem Cytochem 48, 1627–1638. 10.1177/002215540004801206.

75. Perry, G., Kawai, M., Tabaton, M., Onorato, M., Mulvihill, P., Richey, P., Morandi, A., Connolly, J.A., and Gambetti, P. (1991). Neuropil threads of Alzheimer’s disease show a marked alteration of the normal cytoskeleton. J Neurosci 11, 1748–1755. 10.1523/JNEUROSCI.11-06-01748.1991.

76. Ashford, J.W., Soultanian, N.S., Zhang, S.-X., and Geddes, J.W. (1998). Neuropil Threads are Collinear With MAP2 Immunostaining in Neuronal Dendrites of Alzheimer Brain: Journal of Neuropathology and Experimental Neurology 57, 972–978. 10.1097/00005072-199810000-00009.

77. Matusova, Z., Hol, E.M., Pekny, M., Kubista, M., and Valihrach, L. (2023). Reactive astrogliosis in the era of single-cell transcriptomics. Front. Cell. Neurosci. 17, 1173200. 10.3389/fncel.2023.1173200.

78. Samelson, A.J., Ariqat, N., McKetney, J., Rohanitazangi, G., Bravo, C.P., Bose, R., Travaglini, K.J., Lam, V.L., Goodness, D., Dixon, G., et al. (2024). CRISPR screens in iPSC-derived neurons reveal principles of tau proteostasis. bioRxiv, 2023.06.16.545386. 10.1101/2023.06.16.545386.

79. Chaggar, P., Vogel, J.W., Binette, A.P., Thompson, T.B., Strandberg, O., Mattsson-Carlgren, N., Karlsson, L., Stomrud, E., Jbabdi, S., Magon, S., et al. (2025). Personalised regional modelling predicts tau progression in the human brain. PLoS Biol 23, e3003241. 10.1371/journal.pbio.3003241.

80. Vogel, J.W., Iturria-Medina, Y., Strandberg, O.T., Smith, R., Levitis, E., Evans, A.C., Hansson, O., Alzheimer’s Disease Neuroimaging Initiative, and Swedish BioFinder Study (2020). Spread of pathological tau proteins through communicating neurons in human Alzheimer’s disease. Nat Commun 11, 2612. 10.1038/s41467-020-15701-2.

81. Frost, B., and Diamond, M.I. (2010). Prion-like mechanisms in neurodegenerative diseases. Nat Rev Neurosci 11, 155–159. 10.1038/nrn2786.

82. Goedert, M., Clavaguera, F., and Tolnay, M. (2010). The propagation of prion-like protein inclusions in neurodegenerative diseases. Trends in Neurosciences 33, 317–325. 10.1016/j.tins.2010.04.003.

83. Winfree, R.L., Nolan, E., Blennow, K., Zetterberg, H., Gifford, K.A., Pechman, K.R., Schneider, J., Bennett, D.A., Petyuk, V.A., Jefferson, A.L., et al. (2025). Vascular endothelial growth factor receptor-1 (FLT1) interactions with amyloid-beta in Alzheimer’s disease: A putative biomarker of amyloid-induced vascular damage. Neurobiol Aging 147, 141–149. 10.1016/j.neurobiolaging.2024.12.010.

84. Mahoney, E.R., Dumitrescu, L., Moore, A.M., Cambronero, F.E., De Jager, P.L., Koran, M.E.I., Petyuk, V.A., Robinson, R.A.S., Goyal, S., Schneider, J.A., et al. (2021). Brain expression of the vascular endothelial growth factor gene family in cognitive aging and alzheimer’s disease. Mol Psychiatry 26, 888–896. 10.1038/s41380-019-0458-5.

85. Yang, H.-S., Yau, W.-Y.W., Carlyle, B.C., Trombetta, B.A., Zhang, C., Shirzadi, Z., Schultz, A.P., Pruzin, J.J., Fitzpatrick, C.D., Kirn, D.R., et al. (2024). Plasma VEGFA and PGF impact longitudinal tau and cognition in preclinical Alzheimer’s disease. Brain 147, 2158–2168. 10.1093/brain/awae034.

86. Levites, Y., Das, P., Price, R.W., Rochette, M.J., Kostura, L.A., McGowan, E.M., Murphy, M.P., and Golde, T.E. (2006). Anti-Abeta42- and anti-Abeta40-specific mAbs attenuate amyloid deposition in an Alzheimer disease mouse model. J Clin Invest 116, 193–201. 10.1172/JCI25410.

87. Ryu, J.K., Cho, T., Choi, H.B., Wang, Y.T., and McLarnon, J.G. (2009). Microglial VEGF receptor response is an integral chemotactic component in Alzheimer’s disease pathology. J Neurosci 29, 3–13. 10.1523/JNEUROSCI.2888-08.2009.

88. Seto, M., Dumitrescu, L., Mahoney, E.R., Sclafani, A.M., De Jager, P.L., Menon, V., Koran, M.E.I., Robinson, R.A., Ruderfer, D.M., Cox, N.J., et al. (2023). Multi-omic characterization of brain changes in the vascular endothelial growth factor family during aging and Alzheimer’s disease. Neurobiology of Aging 126, 25–33. 10.1016/j.neurobiolaging.2023.01.010.

89. Kisler, K., Nikolakopoulou, A.M., and Zlokovic, B.V. (2021). Microglia have a grip on brain microvasculature. Nat Commun 12, 5290. 10.1038/s41467-021-25595-3.

90. Chen, S., Li, J., Meng, S., He, T., Shi, Z., Wang, C., Wang, Y., Cao, H., Huang, Y., Zhang, Y., et al. (2023). Microglia and macrophages in the neuro-glia-vascular unit: From identity to functions. Neurobiology of Disease 179, 106066. 10.1016/j.nbd.2023.106066.

91. Rymo, S.F., Gerhardt, H., Wolfhagen Sand, F., Lang, R., Uv, A., and Betsholtz, C. (2011). A Two-Way Communication between Microglial Cells and Angiogenic Sprouts Regulates Angiogenesis in Aortic Ring Cultures. PLoS ONE 6, e15846. 10.1371/journal.pone.0015846.

92. Steinman, J., Sun, H.-S., and Feng, Z.-P. (2020). Microvascular Alterations in Alzheimer’s Disease. Front Cell Neurosci 14, 618986. 10.3389/fncel.2020.618986.

93. Iturria-Medina, Y., Sotero, R.C., Toussaint, P.J., Mateos-Pérez, J.M., Evans, A.C., and Alzheimer’s Disease Neuroimaging Initiative (2016). Early role of vascular dysregulation on late-onset Alzheimer’s disease based on multifactorial data-driven analysis. Nat Commun 7, 11934. 10.1038/ncomms11934.

94. Peng, X., Trambaiolli, L.R., Choi, E.Y., Lehman, J.F., Linn, G., Russ, B.E., Schroeder, C.E., Haber, S.N., and Liu, H. (2024). Cross-species striatal hubs: Linking anatomy to resting-state connectivity. NeuroImage 301, 120866. 10.1016/j.neuroimage.2024.120866.

95. 95. Vogel, J.W., Young, A.L., Oxtoby, N.P., Smith, R., Ossenkoppele, R., Strandberg, O.T., La Joie, R., Aksman, L.M., Grothe, M.J., Iturria-Medina, Y., et al. (2021). Four distinct trajectories of tau deposition identified in Alzheimer’s disease. Nat Med 27, 871–881. 10.1038/s41591-021-01309-6.

96. Chu, Y., Hirst, W.D., Federoff, H.J., Harms, A.S., Stoessl, A.J., and Kordower, J.H. (2024). Nigrostriatal tau pathology in parkinsonism and Parkinson’s disease. Brain 147, 444–457. 10.1093/brain/awad388.

97. Duquette, A., Pernègre, C., Veilleux Carpentier, A., and Leclerc, N. (2021). Similarities and Differences in the Pattern of Tau Hyperphosphorylation in Physiological and Pathological Conditions: Impacts on the Elaboration of Therapies to Prevent Tau Pathology. Front. Neurol. 11, 607680. 10.3389/fneur.2020.607680.

98. Mathys, H., Boix, C.A., Akay, L.A., Xia, Z., Davila-Velderrain, J., Ng, A.P., Jiang, X., Abdelhady, G., Galani, K., Mantero, J., et al. (2024). Single-cell multiregion dissection of Alzheimer’s disease. Nature 632, 858–868. 10.1038/s41586-024-07606-7.

99. Ruden, J.B., Dugan, L.L., and Konradi, C. (2021). Parvalbumin interneuron vulnerability and brain disorders. Neuropsychopharmacology 46, 279–287. 10.1038/s41386-020-0778-9.

100. Zhang, N., Hu, B.-W., Li, X.-M., and Huang, H. (2025). Rethinking parvalbumin: From passive marker to active modulator of hippocampal circuits. IBRO Neurosci Rep 19, 760–773. 10.1016/j.ibneur.2025.10.005.

101. Pinna, A., and Colasanti, A. (2021). The Neurometabolic Basis of Mood Instability: The Parvalbumin Interneuron Link-A Systematic Review and Meta-Analysis. Front Pharmacol 12, 689473. 10.3389/fphar.2021.689473.

102. Hijazi, S., Smit, A.B., and Van Kesteren, R.E. (2023). Fast-spiking parvalbumin-positive interneurons in brain physiology and Alzheimer’s disease. Mol Psychiatry 28, 4954–4967. 10.1038/s41380-023-02168-y.

103. Verret, L., Mann, E.O., Hang, G.B., Barth, A.M.I., Cobos, I., Ho, K., Devidze, N., Masliah, E., Kreitzer, A.C., Mody, I., et al. (2012). Inhibitory interneuron deficit links altered network activity and cognitive dysfunction in Alzheimer model. Cell 149, 708–721. 10.1016/j.cell.2012.02.046.

104. Kumar, P., Goettemoeller, A.M., Espinosa-Garcia, C., Tobin, B.R., Tfaily, A., Nelson, R.S., Natu, A., Dammer, E.B., Santiago, J.V., Malepati, S., et al. (2024). Native-state proteomics of Parvalbumin interneurons identifies unique molecular signatures and vulnerabilities to early Alzheimer’s pathology. Nat Commun 15, 2823. 10.1038/s41467-024-47028-7.

105. Hijazi, S., Heistek, T.S., Scheltens, P., Neumann, U., Shimshek, D.R., Mansvelder, H.D., Smit, A.B., and van Kesteren, R.E. (2020). Early restoration of parvalbumin interneuron activity prevents memory loss and network hyperexcitability in a mouse model of Alzheimer’s disease. Mol Psychiatry 25, 3380–3398. 10.1038/s41380-019-0483-4.

106. Lee, K., Holley, S.M., Shobe, J.L., Chong, N.C., Cepeda, C., Levine, M.S., and Masmanidis, S.C. (2017). Parvalbumin Interneurons Modulate Striatal Output and Enhance Performance during Associative Learning. Neuron 93, 1451–1463.e4. 10.1016/j.neuron.2017.02.033.

107. Hodge, R.D., Bakken, T.E., Miller, J.A., Smith, K.A., Barkan, E.R., Graybuck, L.T., Close, J.L., Long, B., Johansen, N., Penn, O., et al. (2019). Conserved cell types with divergent features in human versus mouse cortex. Nature 573, 61–68. 10.1038/s41586-019-1506-7.

108. Aevermann, B., Zhang, Y., Novotny, M., Keshk, M., Bakken, T., Miller, J., Hodge, R., Lelieveldt, B., Lein, E., and Scheuermann, R.H. (2021). A machine learning method for the discovery of minimum marker gene combinations for cell type identification from single-cell RNA sequencing. Genome Res. 31, 1767–1780. 10.1101/gr.275569.121.

109. Wolf, F.A., Angerer, P., and Theis, F.J. (2018). SCANPY: large-scale single-cell gene expression data analysis. Genome Biol 19, 15. 10.1186/s13059-017-1382-0.

110. Virtanen, P., Gommers, R., Oliphant, T.E., Haberland, M., Reddy, T., Cournapeau, D., Burovski, E., Peterson, P., Weckesser, W., Bright, J., et al. (2020). SciPy 1.0: fundamental algorithms for scientific computing in Python. Nat Methods 17, 261–272. 10.1038/s41592-019-0686-2.

111. Mitchel, J., Gao, T., Cole, E., Petukhov, V., and Kharchenko, P.V. (2025). Impact of Segmentation Errors in Analysis of Spatial Transcriptomics Data. Preprint, 10.1101/2025.01.02.631135 https://doi.org/10.1101/2025.01.02.631135.

112. Phan, D., Pradhan, N., and Jankowiak, M. (2019). Composable Effects for Flexible and Accelerated Probabilistic Programming in NumPyro. Preprint at arXiv, 10.48550/ARXIV.1912.11554 https://doi.org/10.48550/ARXIV.1912.11554.

113. Kumar, R., Carroll, C., Hartikainen, A., and Martin, O. (2019). ArviZ a unified library for exploratory analysis of Bayesian models in Python. JOSS 4, 1143. 10.21105/joss.01143.

114. Heumos, L., Ji, Y., May, L., Green, T., Zhang, X., Wu, X., Ostner, J., Peidli, S., Schumacher, A., Hrovatin, K., et al. (2024). Pertpy: an end-to-end framework for perturbation analysis. Preprint, 10.1101/2024.08.04.606516 https://doi.org/10.1101/2024.08.04.606516.

115. He, L., Davila-Velderrain, J., Sumida, T.S., Hafler, D.A., Kellis, M., and Kulminski, A.M. (2021). NEBULA is a fast negative binomial mixed model for differential or co-expression analysis of large-scale multi-subject single-cell data. Communications Biology 4, 629. 10.1038/s42003-021-02146-6.

116. Heumos, L., Schaar, A.C., Lance, C., Litinetskaya, A., Drost, F., Zappia, L., Lücken, M.D., Strobl, D.C., Henao, J., Curion, F., et al. (2023). Best practices for single-cell analysis across modalities. Nat Rev Genet 24, 550–572. 10.1038/s41576-023-00586-w.

117. Janesick, A., Shelansky, R., Gottscho, A.D., Wagner, F., Williams, S.R., Rouault, M., Beliakoff, G., Morrison, C.A., Oliveira, M.F., Sicherman, J.T., et al. (2023). High resolution mapping of the tumor microenvironment using integrated single-cell, spatial and in situ analysis. Nat Commun 14, 8353. 10.1038/s41467-023-43458-x.

118. Marconato, L., Palla, G., Yamauchi, K.A., Virshup, I., Heidari, E., Treis, T., Vierdag, W.-M., Toth, M., Stockhaus, S., Shrestha, R.B., et al. (2025). SpatialData: an open and universal data framework for spatial omics. Nat Methods 22, 58–62. 10.1038/s41592-024-02212-x.

119. van der Walt, S., Schönberger, J.L., Nunez-Iglesias, J., Boulogne, F., Warner, J.D., Yager, N., Gouillart, E., Yu, T., and contributors, the scikit-image (2014). scikit-image: Image processing in Python. 10.48550/ARXIV.1407.6245.

120. Pedregosa, F., Varoquaux, G., Gramfort, A., Michel, V., Thirion, B., Grisel, O., Blondel, M., Prettenhofer, P., Weiss, R., and Dubourg, V. (2011). Scikit-learn: Machine learning in Python. Journal of machine learning research 12, 2825–2830.

121. Leutenegger, S.T., Lopez, M.A., and Edgington, J. (1997). STR: a simple and efficient algorithm for R-tree packing. In Proceedings 13th International Conference on Data Engineering (IEEE Comput. Soc. Press), pp. 497–506. 10.1109/ICDE.1997.582015.

