## Supplemental Information for "The Caudate Nucleus Exhibits Distinct Pathology and Cell Type-Specific Responses Across Alzheimer’s Disease"

**Document S1** Contains Supplemental Figure 1, 2 and 3.

**Table S1** List of genes included in the gene panel of the xenium experiments along with the number of probes for each gene.

**Table S2** Digital pathology measurements for AT8, 6E10, and TDP-43 in caudate and middle temporal gyrus.

**Table S3** Differential abundance analysis results from scCODA.

**Table S4-6** Differential expression analysis results from NEBULA, across global, AT8 and 6E10 CPS respectively. These tables can be found at this DOI: 10.6084/m9.figshare.31043686

**Table S7** CPS measurements along with pathology measurements used to infer CPS for each donor.

**Table S8** Donor cohort demographic and neuropathological metadata.

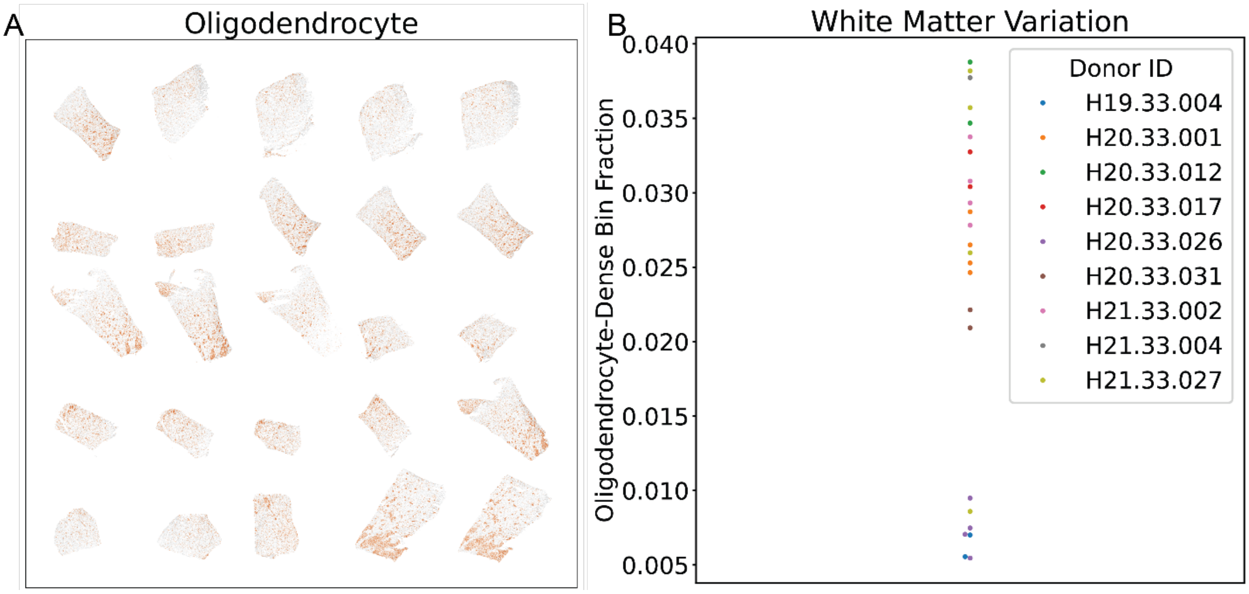


**Supplemental Figure 1. Spatial density analysis of oligodendrocytes**
(A) Scatter plot of cells arranged by their spatial coordinates further arranged into a slab of sections across all donors. Points colored in reddish-brown are oligodendrocytes. B) Swarm plot of sections plotted in (A). Points colored by donor. Sections were converted into in 2500 $\mu m^2$ bins. Oligodendrocyte-dense bins were defined as bins with at least a 10% areal overlap.

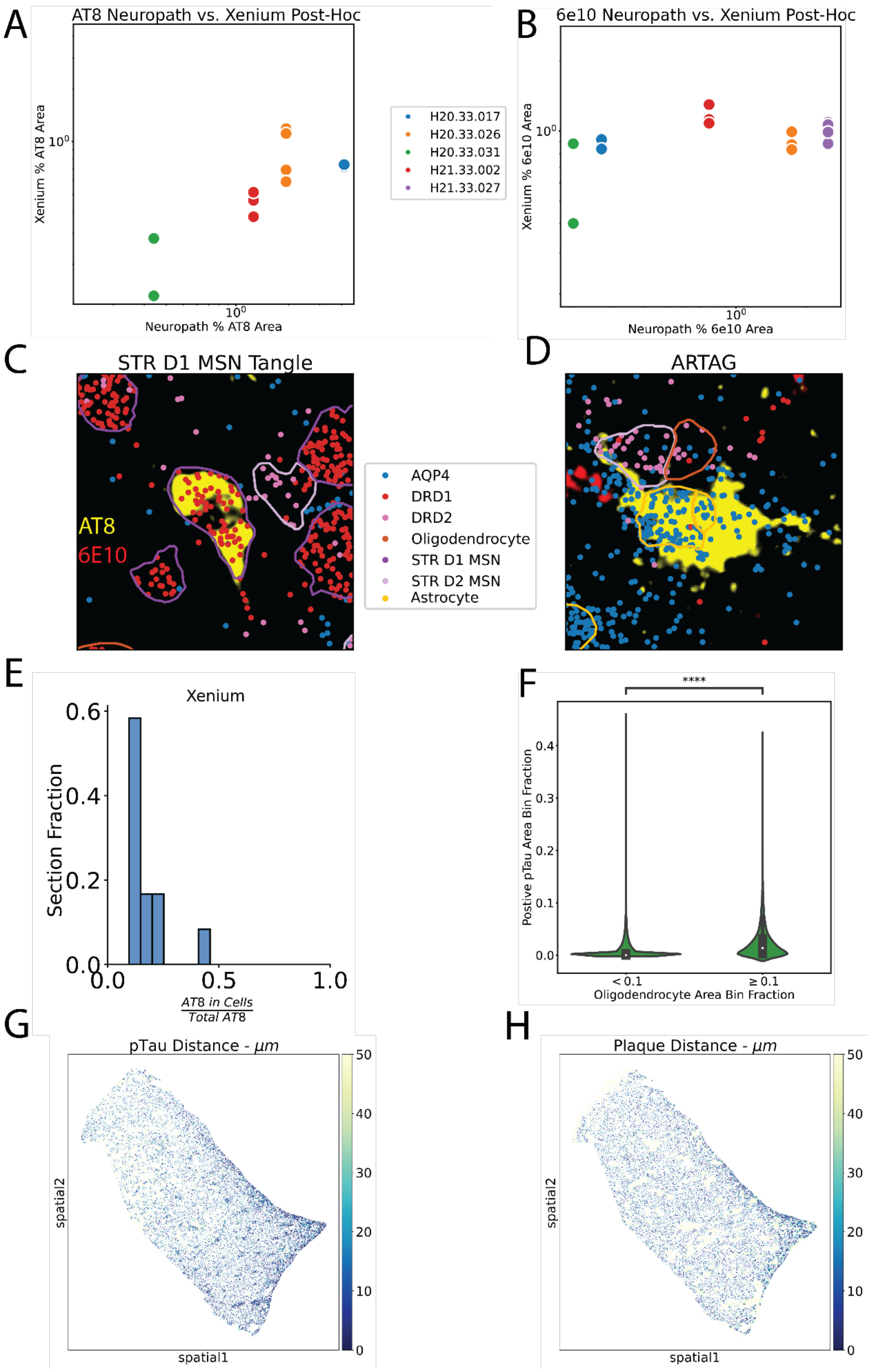
A
AT8 Neuropath vs. Xenium Post-Hoc
Xenium % AT8 Area
Neuropath % AT8 Area
H20.33.017
H20.33.026
H20.33.031
H21.33.002
H21.33.027
B
6e10 Neuropath vs. Xenium Post-Hoc
Xenium % 6e10 Area
Neuropath % 6e10 Area
C
STR D1 MSN Tangle
AT8
6E10
AQP4
DRD1
DRD2
Oligodendrocyte
STR D1 MSN
STR D2 MSN
Astrocyte
D
ARTAG
E
Xenium
Section Fraction
AT8 in Cells / Total AT8
F
****
Postive pTau Area Bin Fraction
< 0.1
≥ 0.1
Oligodendrocyte Area Bin Fraction
G
pTau Distance - μm
spatial2
spatial1
H
Plaque Distance - μm
spatial2
spatial1

**Supplemental Figure 2. Validation  and spatial distribution of post-hoc immunofluorescence.** (A, B) Scatter plots comparing digital neuropathology and Xenium spatial transcriptomics in high pathology load sections. Points colored by donor. (C, D) Post-hoc immunofluorescence images overlaid with transcripts and cell segmentation boundaries in a section from H21.33.002. Segmentation boundaries are colored in yellow. Both panels have AT8 in yellow and 6E10 immunofluorescence intensity. (G) illustrates tangle in D1 MSN neuron. (H) illustrates aging related tau astroglipathy (ARTAG). (E) Histogram of total AT8 signal in cells across high pTau load sections. (F) Violin plot comparing pTau area overlap in 2500 $\mu m^2$ bins in bins with high oligodendrocyte density (overlap $\geq$ 10%) and low oligodendrocyte density (overlap $<$ 10%). ****p < 1e-4 wilocoxon rank sum test. (G, H) Scatter plot of cells arranged by their spatial coordinates in a section from H21.33.002. Points colored by distance to pathology ((G) pTau, (H) A$\beta$ plaques).

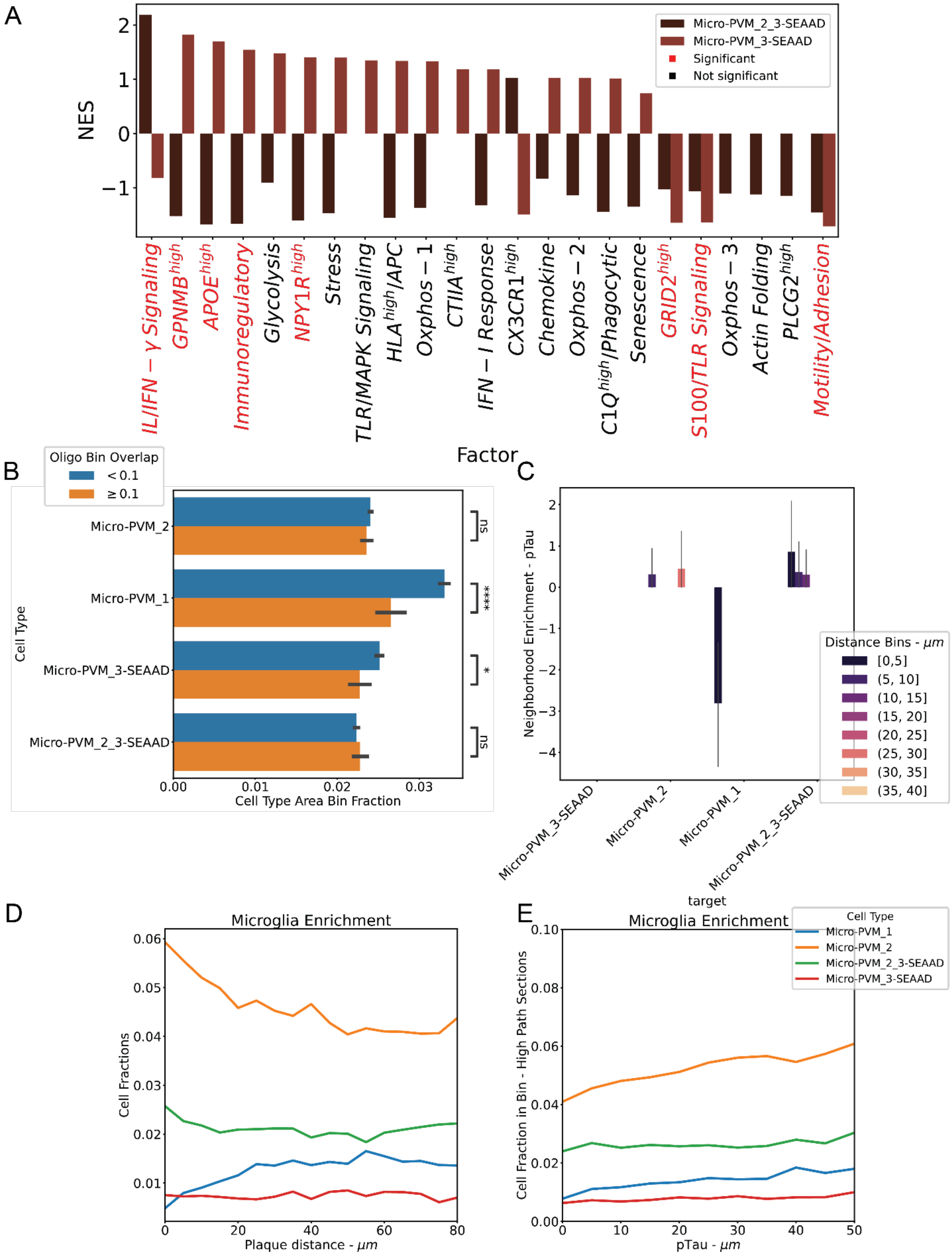


39

40 **Supplemental Figure 3. Further expression and spatial characterization of microglia.**

(A) Bar plot of normalized enrichment score (NES) of marker genes for microglia factors as determined by the *prerank* function from GSEApy[63]. Significance, represented by red ticks, was defined at $p_{adj} < 0.05$. (B) Barplot plot comparing overlap area overlap for different microglia in 2500 $\mu m^2$ bins in bins with high oligodendrocyte density (overlap ≥ 10%) and low oligodendrocyte density (overlap < 10%). ****p < 1e-4 Wilcoxon rank sum test. *p < 0.05 Wilcoxon rank sum test. ns p > 0.05 wilcoxon rank sum. Error bars represent 95% confidence interval across all sections. (C) Bar plot of neighborhood enrichment scores with respect to pTau plaques from spatial transcriptomics. Error bars represent 95% confidence interval across high pTau load sections. (D) Line plot of cellular proportions of 5 $\mu m$ distance bins from A$\beta$ plaques from spatial transcriptomics. Values pooled across all high A$\beta$ load sections. (E) Line plot of cellular proportions of 5 $\mu m$ distance bins from pTau from spatial transcriptomics. Values pooled across all high pTau load sections.
